# RapidAIM: A culture- and metaproteomics-based Rapid Assay of Individual Microbiome responses to drugs

**DOI:** 10.1101/543256

**Authors:** Leyuan Li, Zhibin Ning, Xu Zhang, Janice Mayne, Kai Cheng, Alain Stintzi, Daniel Figeys

## Abstract

**Background:** Human-targeted drugs may exert off-target effects on the gut microbiota. However, our understanding of such effects is limited due to a lack of rapid and scalable assay to comprehensively assess microbiome responses to drugs. Drugs can drastically change the overall microbiome abundance, microbial composition and functions of a gut microbiome. Although we could comprehensively observe these microbiome responses using a series of tests, for the purpose of a drug screening, it is important to decrease the number of analytical tools used.

**Results:** Here, we developed an approach to screen compounds against individual microbiomes *in vitro* using metaproteomics adapted for both absolute bacterial abundances and functional profiling of the microbiome. Our approach was evaluated by testing 43 compounds (including four antibiotics) against five individual microbiomes. The method generated technically highly reproducible readouts, including changes of overall microbiome abundance, microbiome composition and functional pathways. Results show that besides the antibiotics, compounds berberine and ibuprofen inhibited the accumulation of biomass during *in vitro* growth of the microbiome. By comparing genus and species level-biomass contributions, selective antibacterial-like activities were found with 36 of the 39 non-antibiotic compounds. Seven of our compounds led to a global alteration of the metaproteome, with apparent compound-specific patterns of functional responses. The taxonomic distributions of responded proteins varied among drugs, i.e. different drugs affect functions of different members of the microbiome. We also showed that bacterial function can shift in response to drugs without a change in the abundance of the bacteria.

**Conclusions:** Current drug-microbiome interaction studies largely focus on relative microbiome composition and microbial drug metabolism. In contrast, our workflow enables multiple insights into microbiome absolute abundance and functional responses to drugs using metaproteomics as the one-stop screening tool. The workflow is robust, reproducible and quantitative, and is scalable for personalized high-throughput drug screening applications.

## Background

Human-targeted drugs are primarily developed for their effects on the host, and little is known on their effects on the microbiome. Microbiome response to drugs could contribute to off-target drug effect [1]. In addition, the gut microbiome has been linked to gastroenterological, neurologic, respiratory, metabolic, hepatic, and cardiovascular diseases [2]. Therefore, targeting the microbiome could lead to novel therapies [3]. Although the effects of some drugs and compounds on the microbiome have been reported [4], many drug-microbiome interactions are unknown. This is due in part to the extremely high numbers of marketed drugs [5] and compounds in development [6] together with the lack of assays that can rapidly and comprehensively assess the effects of compounds on individual microbiomes. Different *in vitro* approaches have been employed to study drug-microbiome interactions. One strategy involves long term stabilization of the microbiome, as shown in various intestinal microbiome simulators based on continuous flow [7–9]. This approach typically requires a long culture period to stabilize the microbiome (15-20 days), and notable shifts in taxonomic compositions compared with the inoculum have been shown [7, 10]. Moreover, the size and complexity of these culturing systems limit the number of individual microbiomes and drugs that can be examined [9], and thus may not be suitable for high-throughput drug screening purpose. Another strategy is to culture individual bacteria strains isolated from microbiomes. A recent study examined the effects of approved drugs on the biomass of forty individually-cultured bacterial strains in a high-throughput manner [11]. This approach highlighted the importance of biomass in identifying antibacterial-like effects. However, it did not take into account the complexity of a microbial community that could lead to different microbial responses.

Approaches such as optical density measurement [11], flow cytometry [12] and quantitative real-time PCR [13] can be used to compare microbiome biomass. However, these approaches lack insights into drug impact on microbial composition and functions, which are highly related to healthy and disease states. Although we could comprehensively observe microbiome responses by combining multiple tools, for the purpose of initial drug screening, it is important to minimize the number of analytical tools used. There has been no report of an *in vitro* gut microbiome-based drug screening approach that could assess both biomass responses and functional alterations in one test.

The development of meta-omics approaches has allowed rapid and deep measurement of microbiome compositions and functional activities. Genetic approaches such as 16S rDNA and shotgun metagenomics have been regarded as the “gold standard” in microbiome analysis, providing relative quantifications of microbiome membership composition and functional capabilities [14, 15]. However, different microbial members can differ by several orders of magnitude in biomass [16]. Moreover, there is little insight on which microbial traits actually contribute to the functional activities of the microbiome, as functions predicted from 16S rDNA or metagenomics analyses are not necessarily expressed. Studies have shown that gene copy numbers are not representative of protein levels [17]. In addition, RNA expression have limited correlation to the actual protein abundance [18]. In contrast, mass spectrometry (MS)-based metaproteomics technology allows for deep insight into proteome-level information of the microbiome [19, 20], providing quantified protein abundances that estimate the functional activities of the microbiome. Proteins not only provide the biological activities to the microbiome, but also build up a large amount of biomass in microbial cells. Hence, the metaproteomic readouts can also be used to assess the microbiome biomass and analyze community structure [21]. It has been validated that metaproteomics is a good estimator of biomass contributions of microbiome members [22]. Despite its coverage could not compare to that of the genomic sequencing-based technologies, metaproteomics could confidently quantify proteins of the bacterial species that constitute>90% of the total biomass [23], making it sufficient for a fast-pass drug screening application.

Here we report an approach named Rapid Assay of an Individual’s Microbiome (RapidAIM) facing gut microbiome-targeted drug screening, and evaluated the applicability of metaproteomics for insights of microbiome responses to drugs. Briefly, in RapidAIM, individual microbiomes are cultured in a previously optimized culture system for 24 hours, and the samples are then analyzed using a metaproteomics-based analytical approach. A high-throughput equal-volume based protein extraction and digestion workflow was applied to enable absolute biomass assessment along with the functional profiling. To demonstrate the feasibility and performance of the RapidAIM assay, we carried out a proof-of-concept study involving 43 compounds and five individual gut microbiomes. Microbiome responses including changes in biomass, taxon-specific biomass contributions, taxon-specific functional activities, and detailed responses of interested enzymatic pathways can be obtained following the assay.

## Results

### Development and evaluation of RapidAIM

RapidAIM consists of an optimized microbiome culturing method, an equal-volume based protein extraction and digestion workflow and a metaproteomic analysis pipeline **(Figure 1a)**. Briefly, fresh human stool samples are inoculated in 96-well deep-well plates and cultured with drugs for 24 hours. We have previously optimized the culture model and validated that it maintains the composition and taxon-specific functional activities of individual gut microbiomes in 96-well plates [24]. After 24 hours, the cultured microbiomes are prepared for metaproteomic analysis using a microplate-based metaproteomic sample processing workflow **(Supplementary Figure S1)** adapted from our single-tube protocol [25]. The microplate-based workflow consists of bacterial cell purification, cell lysis under ultra-sonication in 8M urea buffer, in-solution tryptic digestion, and desalting. We validated each step of this workflow and found no significant differences in identification efficiency between 96-well plate processing and single-tube processing **(Supplementary Figure S1)**. To compare total biomass, taxon-specific biomass and pathway contributions between samples in a high-throughput assay format, we applied an equal sample volume strategy to our recently developed metaproteomics techniques [20, 26, 27]. To validate the absolute quantification of microbiome abundance by comparing total peptide intensity, an equal volume of samples from a microbiome dilution series (simulating different levels of drug effects) was taken for tryptic digestion and LC-MS/MS analysis. Summed peptide intensity in each sample showed good linearity (R^2^ = 0.991, **Figure 1b**) with a standard colorimetric protein assay, showing that the total peptide intensity is a good indicator for microbiome biomass levels. Since drugs could cause drastically change in microbiome abundance, we then evaluated whether biomass differences between wells could cause bias in identified functional and taxonomic compositions. We confirmed that the level of total biomass didn’t bias the composition of functional profiles (**Figure 1c**), protein groups (**Supplementary Figure S2a**), and taxonomic abundances (**Supplementary Figure S2b**).

**Figure 1.**
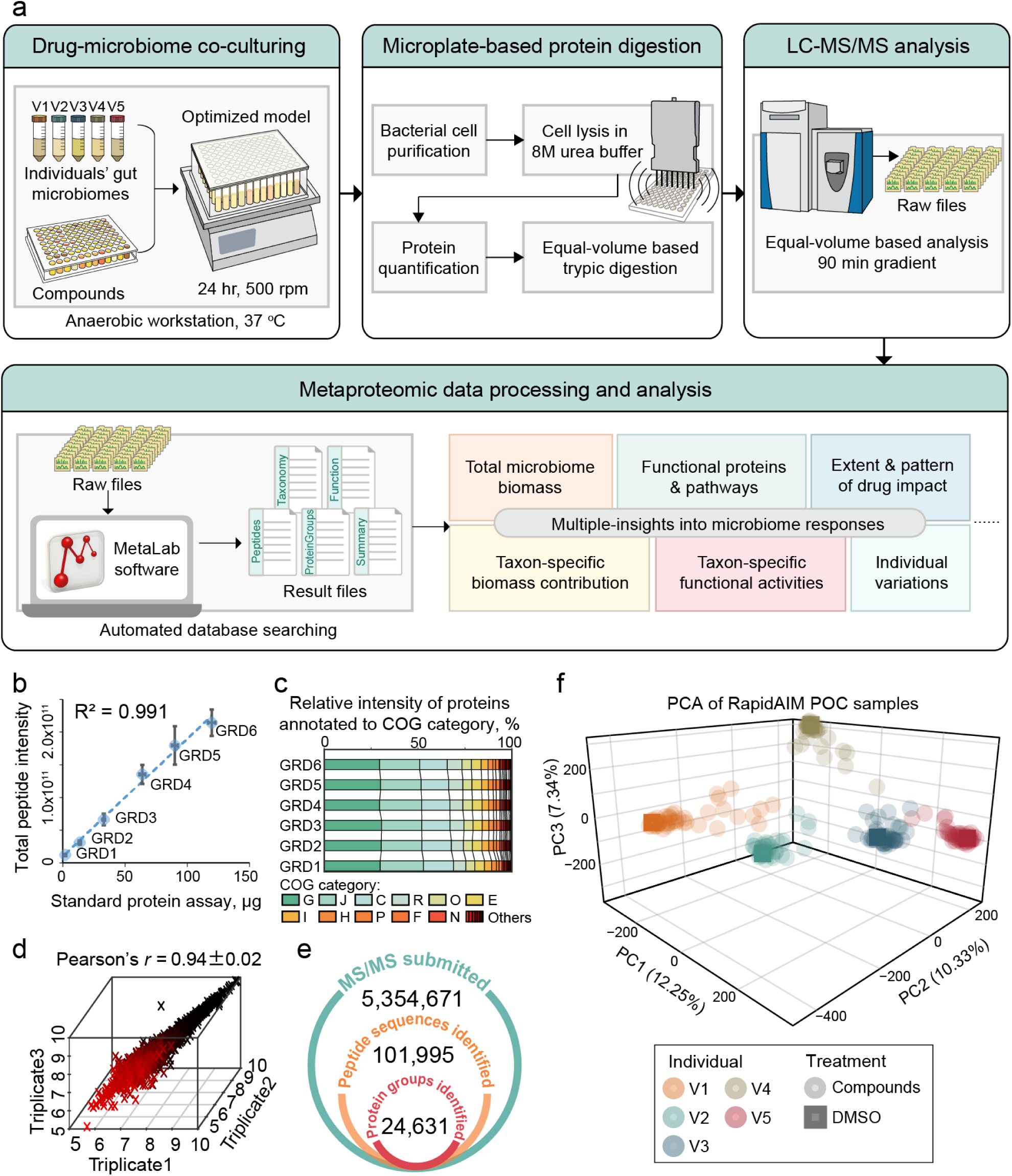
Rapid Assay of Individual Microbiome (RapidAIM) workflow and performance. (**a**) Experimental, analytical, and bioinformatics components of the RapidAIM workflow. Each individual’s gut microbiome samples are cultured with the test compounds in a 96-well deep-well plate at 37° in strict anaerobic conditions for 24 hours followed by high-throughput sample preparation and rapid LC-MS/MS analysis. Peptide and protein identification and quantification, taxonomic profiling, and functional annotation were performed using the automated MetaLab software [27]. (**b**) A series of six dilutions (dilution gradients: GRD1∼6) of a same microbiome sample was tested in triplicate through the equal-volume digestion and equal-volume MS loading protocol; the summed peptide intensity was compared to a set of protein concentration standards provided with DC protein concentration assay and showed good linearity (center points and error bars represent mean ± SD). (**c**) Stacked bars of clusters of orthologous groups (COG) category levels across the six concentrations showing no bias at the functional quantifications. (**d**) Analysis of three technical replicates in RapidAIM showing high Pearson’s correlation. Numbers of MS/MS submitted, peptide sequence and protein group identifications in the POC dataset. PCA based on LFQ intensities of protein groups for all POC samples.

### RapidAIM: Proof-of-concept study

We conducted a proof-of-concept (POC) study on the use of RapidAIM to characterize drug-microbiome interactions. We selected 43 compounds that have been previously suggested to impact, interact with, or be metabolized by the gut microbiome (**Supplementary Table S1)**. Thirty-seven of these compounds are FDA-approved drugs; four are antibiotics, and the others include nonsteroidal anti-inflammatory drugs (NSAIDs), anti-diabetic drugs, aminosalicylate, and statins, etc. Each compound, at a dose corresponding to the assumed maximal fecal concentration of its daily dose, was added to five wells of 96-well plates containing 1 ml culture medium in each well. The drug solvent, dimethyl sulfoxide (DMSO), was used as the negative control. Then, each of the five wells for each compound was inoculated with a different fecal microbiome from healthy human volunteers. Following 24 hours of culturing, the samples were processed through the microplate-based workflow **(Figure S1)** and were subjected to a 90 min gradient-based rapid LC-MS/MS analysis. Using our automated metaproteomic data analysis software MetaLab [27], 101,995 peptide sequences corresponding to 24,631 protein groups were quantified across all samples with a false discovery rate (FDR) threshold of 1% **(Figure 1e)**. The average MS/MS identification rate was 32.4 ± 8.8% (mean ± SD); an average of 15,017 ± 3,654 unique peptides and 6,684 ± 998 protein groups were identified per sample. To provide a global overview of the microbiome responses, a PCA was performed based on label-free quantification (LFQ) intensities of protein groups **(Figure 1f)**. As expected, the samples clustered based on the original microbiome source and not based on drug treatment. Within each individual microbiome group, a number of drug-treated samples clustered closely to their control while several drug-treated samples clearly separated from the non-treated control.

We next evaluated the robustness and reproducibility of the method by culturing one microbiome with drugs in technical triplicates. Cultured triplicates yielded high Pearson’s *r* for LFQ protein group intensities **(Figure 1d**). Hierarchical clustering based on Pearson’s *r* of LFQ protein group intensities between samples showed that with the exception of several compounds which clustered closely with DMSO, cultured triplicates were clustered together **(Supplementary Figure S3a)**. Moreover, total biomass, functional enzymes, and species biomass contributions were highly reproducible between triplicates as shown in **Supplementary Figure S3b-d**.

### Effects of compounds on microbiome abundance and composition

We examined the effect of the 43 compounds on the overall abundance (biomass) of each individual microbiome by comparing the total peptide intensity **(Figure 2a)**. As expected, the antibiotics greatly reduced total microbial biomass in most individual microbiomes (with one exception of increased microbiome abundance in response to rifaximin, further examination is shown in **Supplementary Figure S4**). Closely clustered with these antibiotics, berberine and ibuprofen also inhibited the biomass of all individual microbiomes.

**Figure 2.**
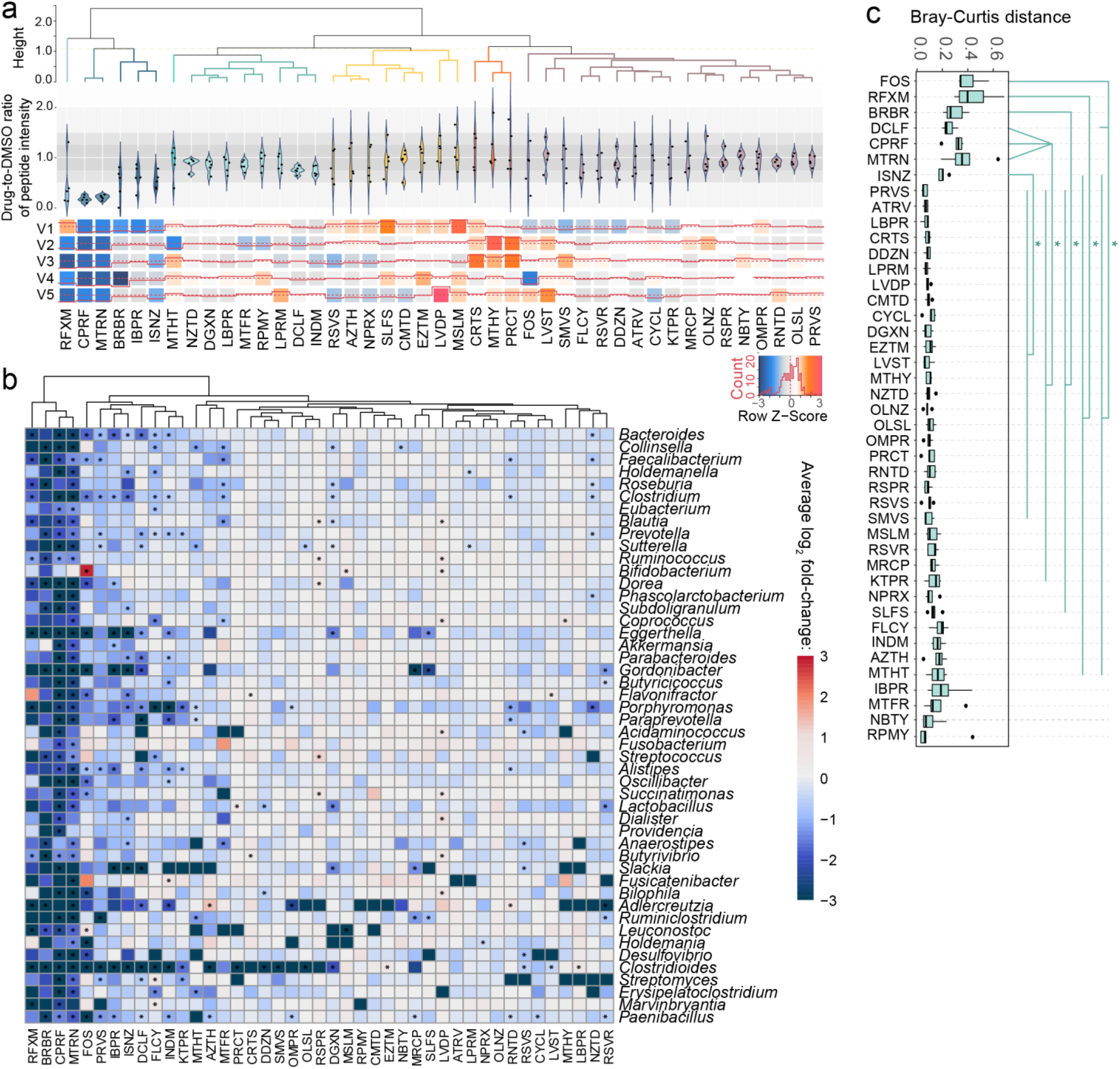
Response of microbiome abundance and composition to compounds. (**a**) Biomass responses of individuals’ microbiomes to compounds relative to DMSO control. Ratio of peptide intensity between compound and DMSO control samples was calculated for each individual microbiome. (**b**) Log2 fold-change of absolute abundance at the genus level in response to each drug compared with the DMSO control. Genera that existed in ≥80% of the volunteers are shown. Star (*) indicate significantly changed bacterial abundance by Wilcoxon test, *p* <0.05. (**c**) Bray-Curtis distance of genus-level composition between drug-treated microbiomes and the corresponding DMSO control samples. Heatmap colors are generated with average of log2-fold changes among the five individual samples. Statistical significance was calculated by pairwise Wilcoxon test (FDR-adjusted *p* < 0.05). Box spans interquartile range (25th to 75th percentile) and line within box denotes median. For full compound names, see abbreviation list in **Supplementary Table S1**.

We next explored the effects of drugs on the microbiome composition based on bacterial biomass contributions. To evaluate the overall shift of the microbiome, Bray-Curtis distance [28, 29] between drug-treated and DMSO control microbiome indicated that fructooligosaccharide (FOS), rifaximin, berberine, diclofenac, ciprofloxacin, metronidazole and isoniazid significantly shifted the microbiome (pairwise Wilcoxon test, FDR-adjusted *p* < 0.05; **Figure 2c**).

Our metaproteomic dataset allowed us to further examine the response of bacterial absolute abundance by comparing summed peptide intensities of each taxon (**Figure 2b**). As expected, the broad-spectrum antibiotics rifaximin, ciprofloxacin and metronidazole significantly inhibited the absolute abundance of a high number of bacterial genera (Wilcoxon test, *p* <0.05). Non-antibiotic compounds, such as berberine, FOS, pravastatin, ibuprofen, diclofencac, flucycosine, and indomethacin also showed significant decrease in the abundances of over ten genera. In addition, selective antibacterial activities were found with 35 out of the other 39 compounds at the genus level. Interestingly, it is clear in **Figure 2b** that while several genera were inhibited, the absolute intensity of *Bifidobacterium* was significantly increased by fructose-oligosaccharides (FOS). Compared to the absolute abundance, the relative abundance provided a different insight into microbiome composition changes (**Supplementary Figure S5).** For example, as opposed to the finding that ciprofloxacin and metronidazole inhibited the biomass of most genera (Figure 2b), they significantly increased the relative abundance of genera *Bifidobacterium, Ruminococcus, Butyrivibrio, Paenibacillus*, etc. Several genera including *Bifidobacterium, Collinsella, Fusobacterium, Butyrivibrio* and *Leuconostoc* were significantly increased in their relative abundances by FOS. Interestingly, members of the Actinobacteria phyla, including *Eggerthella, Gordonibacter, Slackia*, and *Adlercreutzia* were more susceptible to drugs compared to most other genera. Moreover, at the species level, we found that 36 of the 43 compounds significantly affected the biomass of at least one bacterial species (one-sided Wilcoxon rank sum test, FDR-adjusted *p* < 0.05; **Supplementary Table S2**). To this end, RapidAIM allowed for the assessment of changes in both absolute and relative abundances of microbes in response to the compounds.

### Gut microbiome functions in response to compounds

The Bray-Curtis distance of protein group profiles showed that all the four antibiotics, as well as FOS, berberine and diclofenac significantly altered the microbiome functions (**Figure 3a**). These functional alterations likely stemmed from changes in taxonomic composition as revealed by the genus-level Bray-Curtis distance analysis (**Figure 2c)**. We next analyzed the protein group intensities by partial least square discriminant analysis (PLS-DA) to determine whether metaproteomic profiles could be used to discriminate between the DMSO-control and each of the drug-treated microbiomes. In agreement with the Bray-Curtis analysis results, PLS-DA interpretation identified drug-specific metaproteomic patterns associated with seven of our compounds, including the four antibiotics, FOS, berberine and diclofenac (**Supplementary Figure S6**). Hence, hereafter we named these seven compounds as class I compounds, whereas others were named class II compounds. To gain a better understanding of the global effects of class I compounds on the gut microbiome, we applied an unsupervised non-linear dimensionality reduction algorithm, t-distributed stochastic neighbor embedding [30], to visualize this subgroup of metaproteomic data based on protein group abundances (**Figure 3b**). Class I compounds led to a global alteration of the metaproteome, with apparent compound-specific patterns. We next examined the drug impacts on the abundance of functional proteins according to clusters of orthologous groups (COG) of proteins. We identified 535 COGs significantly decreased by at least one drug treatment; 15 of these COGs were decreased by ≥ 10 compounds (**Supplementary Figure S7**). Diclofenac and FOS were the only two compounds that significantly increased COGs (55 and 81 COGs, respectively). Enrichment analysis based on these significantly altered COGs shows that COG categories found to be enriched were responsive to 13 of our compounds (**Figure 3c**), six of those were class I compounds. Interestingly, the non-antibiotic NSAID diclofenac increased the abundance of several COG categories (**Figure 3c**). By mapping these significantly increased proteins from these COG categories against the string database, we found that these altered proteins are functionally interconnected (**Supplementary Figure S8**). Interestingly, one of the proteins that were highly connected in the string network, COG0176 – transaldolase (1.76 ± 0.21 fold-change), is involved in the biosynthesis of ansamycins, bacterial secondary metabolites that have antibiotic activities [31].

**Figure 3.**
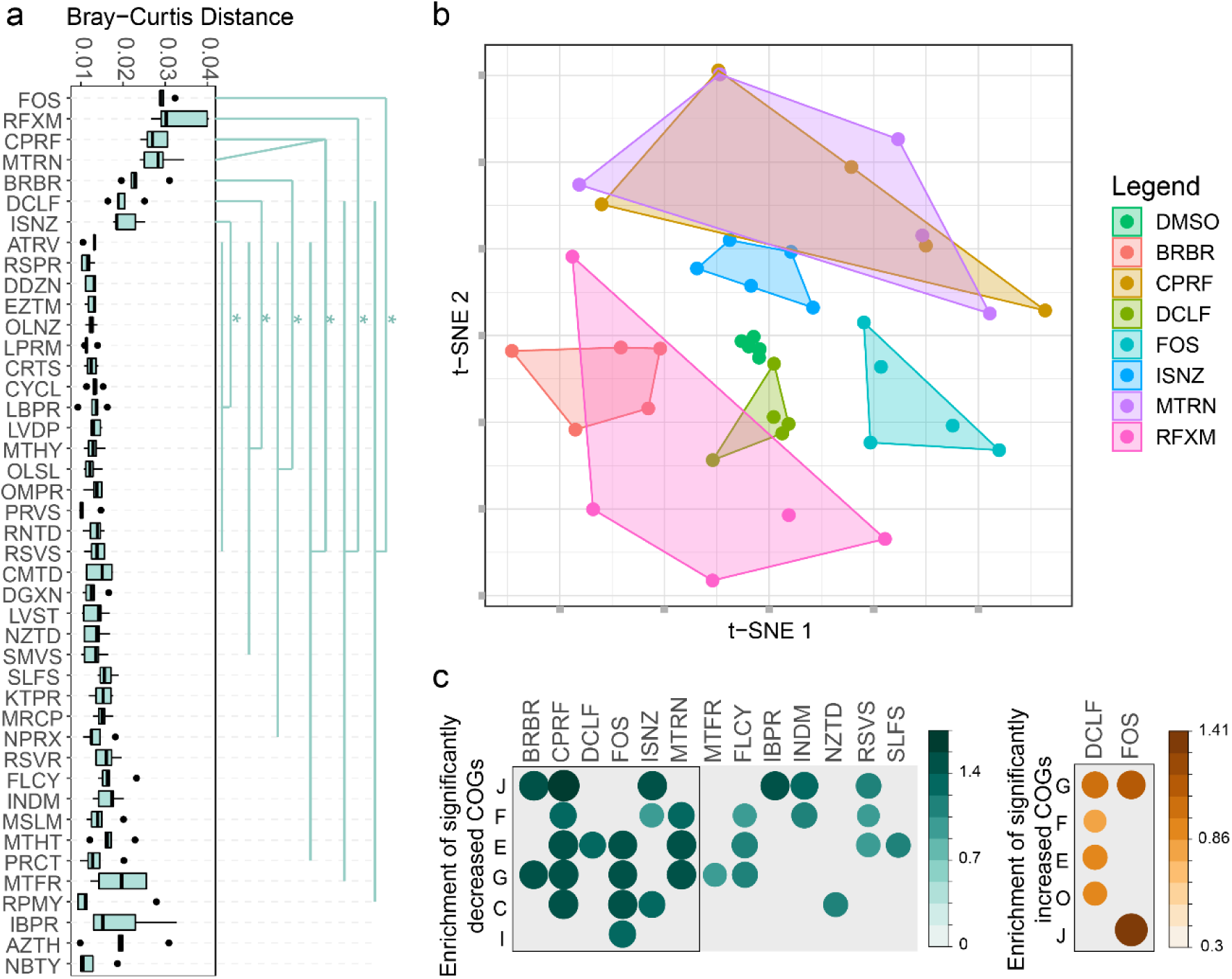
Effect of compounds on metaproteomic profiles of the microbiome. (**a**) Bray-Curtis distance of protein groups between drug-treated microbiomes and the corresponding DMSO control samples. Statistical significance was calculated by pairwise Wilcoxon test (FDR-adjusted *p* < 0.05). (**b**) Unsupervised dimensionality reduction analysis suggesting three different classes of compound effects. (**c**) Enrichment analysis of all significantly different COGs in the POC dataset. Significantly altered COGs with a *p*-value cutoff of 0.05 are shown.

### Taxon-specific functional responses to class I compounds

We next performed a taxonomic analysis of the functional responses to diclofenac, FOS, ciprofloxacin, and berberine, which represent four different types of compounds (NSAID, oligosaccharide, antibiotics, anti-diabetes) in the class I. Protein groups with VIP scores >1 (thereafter defined as differential proteins) were extracted from each model, and were annotated with their taxonomic and COG information. The taxonomic distributions of the differential proteins varied among drugs (**Figure 4a)**. Moreover, mapping of the differential proteins to phyla-specific pathways revealed phyla-specific responses, as shown for berberine in **Supplementary Figure S9.** In agreement with **Figure 4a**, a higher proportion of down-regulated than up-regulated pathways were identified in Firmicutes and Actinobacteria, while the opposite pattern was observed in Bacteroidetes, Proteobacteria and Verrucomicrobia. In some cases, the phylum-specific responses included up-regulation and down-regulation of different proteins within the same pathway (black lines, **Supplementary Figure S9**). For example, we observed this pattern in fatty acid, carbohydrate, and nucleotide metabolism pathways in Firmicutes. Genus-level analysis revealed genus-specific responses to berberine (**Figure 4b**). In most genera, the genus-specific responses correlated with the overall abundance of the corresponding genus (**Figure 4b**, right panel). Nevertheless, some genera showed functional shifts in response to berberine without changes in overall abundance. For example, *Bifidobacteria, Roseburia, Eubacterium, Clostridium, Ruminococcus, Blautia*, and *Subdoligranulum* exhibited down-regulation of proteins in various COG categories but no changes in biomass were observed.

**Figure 4.**
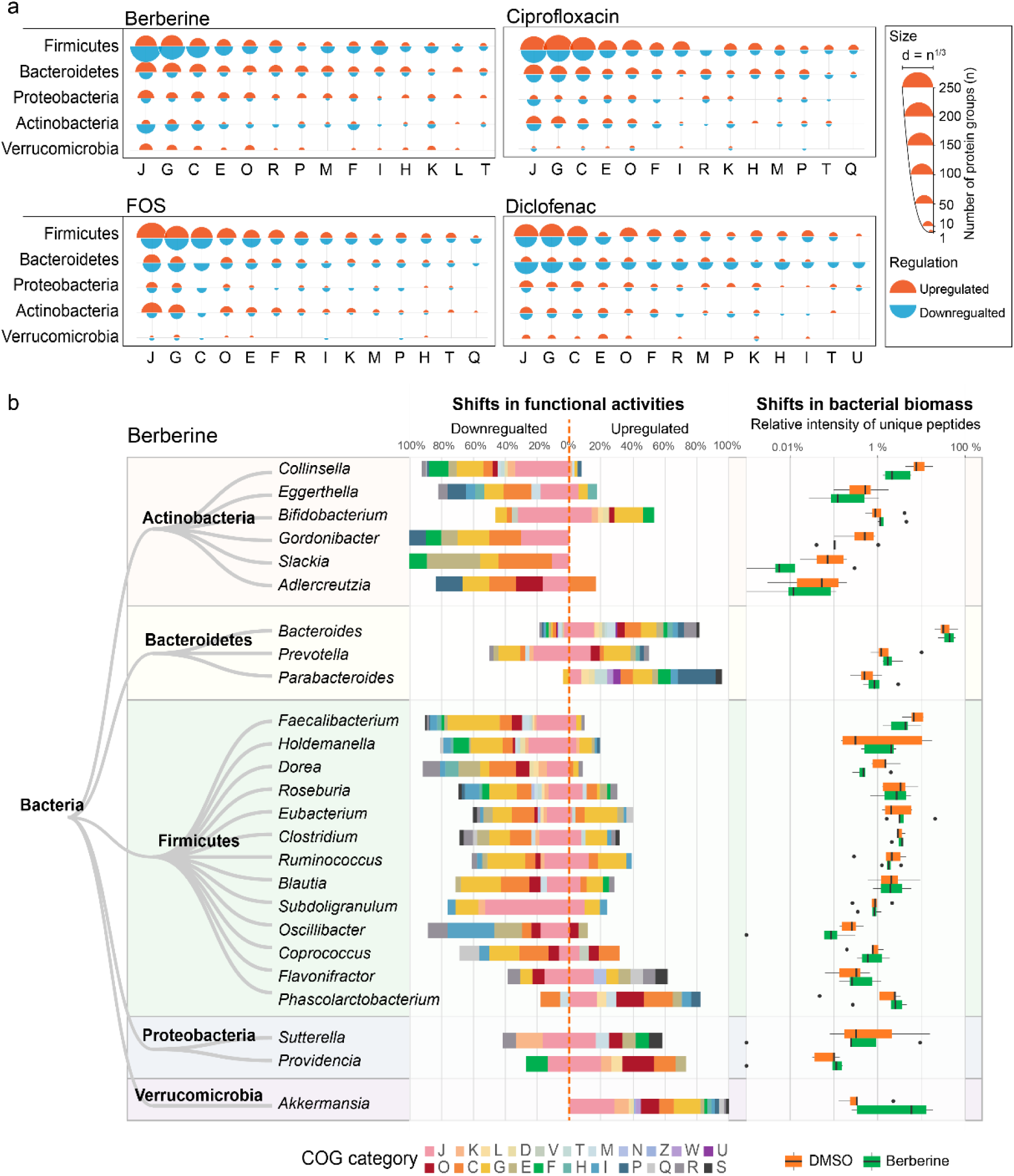
Global functional effects of berberine, ciprofloxacin, FOS and diclofenac. (**a**) Taxon-function distribution of protein groups responding to berberine, FOS, ciprofloxacin and diclofenac. Responding protein groups were selected by PLS-DA based on ComBat-corrected data. The semicircle diameter represents the number of PLS-DA VIP>1 protein groups corresponding to each phyla-COG category pair. (**b**) Genus-level shifts in functional activities in response to berberine and the alterations in biomass of the corresponding genera. Functional shifts (differential protein groups) were identified by PLS-DA. For each genus, the percentages of the total numbers of up- and down-regulated protein groups corresponding to each COG category are shown. Shifts in bacterial biomass in the five microbiomes are shown in box plots with the boxes spanning interquartile range (25th to 75th percentile), and the vertical lines denoting the median for each genus.

### Enzymatic pathways in response to class I compounds

Next we examined the ability of RapidAIM in observing detailed enzymatic pathways of interest. As examples, we show the effects of FOS and ciprofloxacin at the enzymatic pathway level. Protein groups to were annotated to KEGG (Kyoto Encyclopedia of Genes and Genomes) enzymes and were mapped against the KEGG pathway database. **Figure 5a** shows that FOS increased enzymes responsible for fructan and sucrose uptake, as well as enzymes for conversion of D-fructose into D-fructose-1-phosphate, D-mannose-6-phosephate and β-D-fructose-6-phosphate. FOS also affected enzymes involved in the interconversion between glutamine, glutamate and GABA (molecules involved in gut-brain communication). In addition, enzymes involved in sulphide accumulation were affected, including decrease of dissimilatory sulfite reductase (EC 1.8.99.5) and increase of cysteine synthase (EC 2.5.1.47).

**Figure 5.**
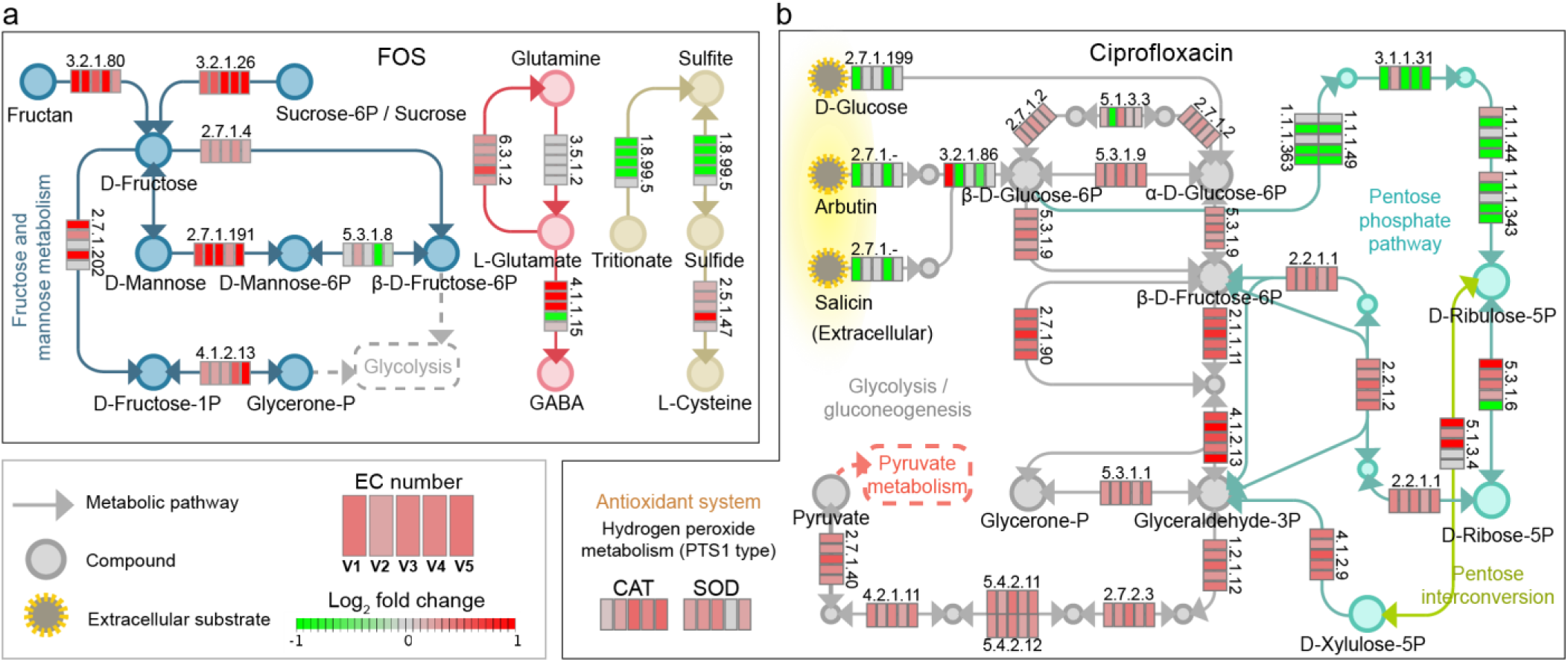
Response of enzymatic pathways to drug treatment. (a) Effect of FOS treatment on enzymes involved in fructose and mannose metabolism, GABA production and sulfide metabolism pathways. (b) effect of ciprofloxacin treatment on enzymes involved in the glycolysis/gluconeogenesis and pentose phosphate pathway. The five blocks of each enzyme represent the five individual microbiomes. Colors in the blocks represent differences between normalized KEGG enzyme intensities with drug treatment versus DMSO (log2-transformation of the original intensity followed by a quotient normalization (x/mean)).

Ciprofloxacin significantly altered the levels of enzymes involved in glycolysis/glycogenesis and pentose phosphate pathways **(Figure 5b)**. The majority of enzymes involved in glycolysis were significantly increased by ciprofloxacin. Ciprofloxacin down-regulated enzymes (ECs 1.1.1.49/1.1.1.363, 3.1.1.31, 1.1.1.44/1.1.1.343) involved in synthesis of ribulose-5-phosphate, which can be isomerized to ribose 5-phosphate for nucleotide biosynthesis [32]. Moreover, the levels of antioxidant enzymes superoxide dismutase (SOD) and catalase (CAT) were increased, suggesting that ciprofloxacin induces oxidative stress in gut bacteria.

### Gut microbiome functions altered by class II compounds

Class II compounds, in contrast to class I compounds, did not cause a global shift in the five individual microbiomes (an example is given by indomethacin, **Figure 6a**). However, **Figure 6a** as well as the Bray-Curtis analyses (**Figure 2c** and **3a**) suggest that there were could be individual variabilities in the extent of drug response. We show that if analyzed on an individual sample basis, significant individualized functional effects could be revealed (**Figure 6b and c**), suggesting high sensitivity of the RapidAIM assay in its application to personalized drug screenings. For example, we identified 303 significantly altered protein groups in cultured replicates of a single indomethacin-treated microbiome (V1). Taxon-function coupled enrichment analysis showed that down-regulated functions were highly enriched in the genus *Bacteroides*, while up-regulated functions were mostly enriched in the order Enterobacterales (**Figure 6d and e**). The up-regulated functions of Enterobacterales included COG0459 chaperonin GroEL (HSP60 family) and COG0234 co-chaperonin GroES (HSP10) (**Figure 6e**).

**Figure 6.**
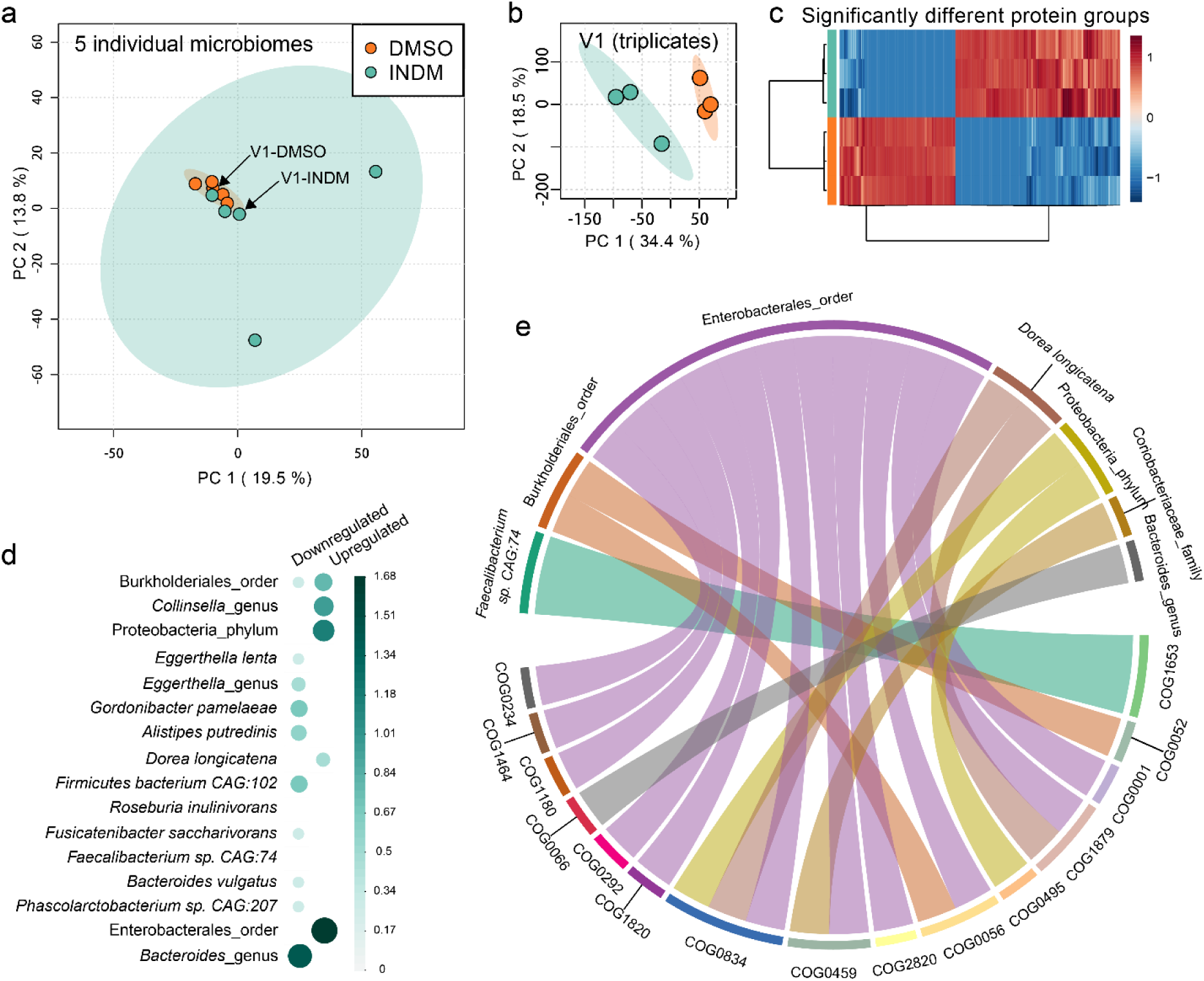
Individual functional responses to indomethacin. (**a**) When visualizing several individual microbiome responses with PCA (based on LFQ intensities of protein groups), inter-individual variability can be greater than drug-induced functional shifts. (**b**) PCA clearly differentiated the response of microbiome V1 treated in triplicates using RapidAIM. (**c**) 303 significant protein group responders were found by *t*-test (FDR-adjusted *p*<0.05). (**d**) Taxon enrichment analysis based on the differential protein groups, (*p*-adjusted=0.05). (**e**) Taxon-function coupled enrichment analysis of up-regulated protein groups.

## Discussion

In the present study, we developed an approach named Rapid Assay of Individual Microbiome (RapidAIM) to evaluate the effects of xenobiotics on individual microbiomes. The range of xenobiotics that reach the intestine and may interact with the gut microbiome is massive and expanding. These xenobiotics include antibiotics and other pharmaceuticals, phytochemicals, polysaccharides, food additives and many other compounds. With the exception of antibiotics, we remain surprisingly ignorant on the extent to which these compounds affect the functions of the gut microbiome. This understanding was limited by lacking an efficient and scalable approach that could maximally obtain insights into microbiome responses while minimizing the number of analytical tools being used.

Here we describe an approach which enables the exploration of drug-microbiome interactions using an optimized *in vitro* culturing model and a metaproteomic approach. We have achieved the maintenance of the representativeness of the initial individual microbiome [24]. Besides, for an *in vitro* culturing simulating the *in vivo* microbiome, it is important to note that the population of gut bacteria in the human body is highly dynamic. It has been estimated that there are ∼0.9·10^11^ bacteria/g wet stool and a total of ∼3.8·10^13^ bacteria in the colon. Approximately 200 g wet daily stool would be excreted [33], leading to a dramatic decrease of the bacterial number in the gut; on the other hand, new bacterial biomass starts growing on nutrients passing through the gut. Current technologies examining the effect of xenobiotic stimulation are usually based on microbiome stabilized after over two weeks of culturing. However, at the stable phase of microbiome growth, the ecosystem has reached its carrying capacity (stable population size), limiting possible observations such as drug effect on the biomass. In our studies, we have previously validated that our composition of gut microbiome is well-maintained along the growth curve [24], so we were able to observe drug responses of growing gut microbiomes by adding the compounds at the initial inoculation stage. Subsequently, combined with our quantitative metaproteomics approach based on equal-sample volume digestion, we were able to observe the drug responses of overall microbiome abundance and taxon-specific biomass contributions.

Here we address the importance of absolute quantification of microbiome biomass contributions. Knowing the change of total microbiome biomass would be helpful to assess the antibiotic-like activity of a compound. As our results clearly showed that the tested antibiotics inhibited the accumulation of microbiome biomass, we found that non-antibiotic compounds ibuprofen and berberine also showed inhibitory effects. Ibuprofen has been frequently used as a safe medication. A study based on relative abundance discussed that ibuprofen had less aggressive effects on the gut microbiome compared to some other NSAIDs [34]. However, in our study, ibuprofen significantly inhibited the overall microbiome biomass through suppressing common gut commensals such as *Bacteroides, Clostridium, Dorea, Eggerthella, Akkermansia*, etc. In terms of berberine, previous studies suggested that berberine has positive effect on beneficial gut microbes, e.g. selectively enriched a few putative short-chain fatty acid producing bacteria [35], and increased the relative abundance of *Akkermansia* spp [36]. However, under the background of an overall inhibition, an enrichment of a taxon (increase in relative abundance) does not necessarily relate to its outgrowth. As another example, our result showed that ciprofloxacin, metronidazole and FOS significantly enriched *Bifidobacterium* (**Supplementary Figure S5**). The fact that *Bifidobacterium* has a certain resistance to ciprofloxacin and metronidazole [37] could contribute to its higher adaptability over other genera. However, we didn’t find evidence of increase in its absolute abundance in presence of ciprofloxacin and metronidazole with RapidAIM. On the contrary, FOS, which could be utilized as the carbon source of *Bifidobacterium* [38, 39], significant increased both absolute and relative abundances of this genus in our study.

We showed that the RapidAIM assay yielded insights into functional responses at multiple levels. Using PLS-DA, we found that berberine, FOS, metronidazole, isoniazid, ciprofloxacin, diclofenac, and rifaximin consistently shifted the metaproteome of the individual gut microbiomes. By annotating the altered proteins at taxonomy, function and pathway levels, we revealed the actions of the different drugs on the microbiome. For example, FOS treatment elevated enzymes involved in fructan and sucrose uptake, as well as enzymes involved in the interconversion among glutamine, glutamate and GABA, which are associated with microbiome communication via the gut-brain axis [40]. In agreement, a study has shown that FOS administration increased GABA receptor genes in mice, and further exhibited both antidepressant and anxiolytic effects [41]. FOS also decreased proteins involved in sulfide generation, suggesting decreased sulfide accumulation in the microbiome. This observation is in agreement with *in vivo* studies showing that FOS treatment decreased the concentration of fecal H_2_S [42–44]. Ciprofloxacin treatment increased enzymes SOD and CAT, which was in agreement with several reports indicating that ciprofloxacin triggers oxidative stress in several bacteria [45–47]. With berberine treatment, we showed that taxon-specific functional shifts can occur either with or without a change in the taxon’s biomass. This observation highlights the strength of our workflow which enables quantitative metaproteomic profiling of the microbiome. Indeed, current classical sequencing-based approaches (16S rDNA or metagenomics sequencing), which generate relative abundances, would not detect these types of changes. Finally, we showed that although a compound may not show global impacts across the five tested microbiomes, it could result in significant alterations on a single microbiome basis. The example given by Indomethacin showed that the order Enterobacterales were enriched with increased chaperonin GroEL (HSP60 family) and co-chaperonin GroES (HSP10) (**Figure 6e**), which have been implicated in infection and diseases pathology [48].

Our workflow still exhibits certain limitations. In particular, MS analysis is a time-consuming process. To this end, a fast-pass screening process such as tandem mass tags (TMT)[49, 50] could be used to multiplex multiple microbiome samples in one MS analysis. Furthermore, our workflow only measures the direct effects of compounds on the microbiome. In its current implementation, it does not take into account the host effect on the microbiome and/or the effects of drug metabolites produced by the host. Future efforts could be aimed at incorporating co-culture of host cells/tissue and gut bacteria [51–53] into a high-throughput drug screening process for achieving more comprehensive insights on host-drug-microbiome interaction. Metaproteomics is a tool that is orthogonal to other omics technologies [17], hence for the need of deeper investigations, RapidAIM could also be coupled with techniques such as metagenomics or metabolomics for a multiple dimension view of the microbiome interaction with drugs.

## Conclusion

To date, the field of drug-microbiome interactions largely focuses on relative microbiome composition and microbial drug metabolism, with a limited understanding of the effects of pharmaceuticals on the absolute abundance and the function of the gut microbiome. A better understanding of these interactions is essential given that the drug effects on the microbiome biomass and functions may have important health consequences. Our workflow enabled the insights into both absolute abundances and functional responses of the gut microbiome to drugs using metaproteomics as the single analytical tool. We have shown that our workflow is robust, reproducible and quantitative, and is easily adaptable for high-throughput drug screening applications.

## Methods

### Stool sample preparation

The Research Ethics Board protocol (# 20160585-01H) for stool sample collection was approved by the Ottawa Health Science Network Research Ethics Board at the Ottawa Hospital. Stool samples were obtained from 5 healthy volunteers (age range 27 - 36 years; 3 males and 2 females). Exclusion criteria were: IBS, IBD, or diabetes diagnosis; antibiotic use or gastroenteritis episode in the last 3 months; use of pro-/pre-biotic, laxative, or anti-diarrheal drugs in the last month; or pregnancy. All volunteers were provided with a stool collection kit, which included a 50 ml Falcon tube containing 15 ml of sterile phosphate-buffered saline (PBS) pre-reduced with 0.1% (w/v) L-cysteine hydrochloride, a 2.5 ml sterile sampling spoon (Bel-Art, United States), plastic wrap, gloves and disposal bags. Briefly, each volunteer placed the plastic wrap over a toilet to prevent the stool from contacting water, collected ∼3 g of stool with the sampling spoon, and dropped the spoon into the prepared 50 ml tube. The sample was immediately weighed by a researcher and transferred into an anaerobic workstation (5% H_2_, 5% CO_2_, and 90% N_2_ at 37°C), where the tube was uncapped to remove O_2_ before homogenization with a vortex mixer. Then the homogenate was filtered using sterile gauzes to remove large particles and obtain the microbiome inoculum.

### Culturing of microbiomes and drug treatments

Each microbiome inoculum was immediately inoculated at a concentration of 2% (w/v) into a 96-well deep well plate containing 1 ml culture medium and a compound dissolved in 5 µl DMSO (or 5 µl DMSO as the control) in each well. The culture medium contained 2.0 g L^−1^ peptone water, 2.0 g L^−1^ yeast extract, 0.5 g L^−1^ L-cysteine hydrochloride, 2 mL L^−1^ Tween 80, 5 mg L^−1^ hemin, 10 μL L^−1^ vitamin K1, 1.0 g L^−1^ NaCl, 0.4 g L^−1^ K_2_HPO_4_, 0.4 g L^−1^ KH_2_PO_4_, 0.1 g L^−1^ MgSO_4_·7H_2_O, 0.1 g L^−1^ CaCl_2_·2H_2_O, 4.0 g L^−1^ NaHCO_3_, 4.0 g L^−1^ porcine gastric mucin (cat# M1778, Sigma-Aldrich), 0.25 g L^−1^ sodium cholate and 0.25 g L^−1^ sodium chenodeoxycholate. The culture medium was sterile and had been pre-reduced overnight in an anaerobic workstation. Concentration of each compound was determined based on the assumption that maximal oral dosage of the drug distributed in 200 g average weight of the colon contents. However, several compounds (i.e. cimetidine, ciprofloxacin, flucytosine, mesalamine, metformin, metronidazole, naproxen-sodium, paracetamol, rifaximin, sodium butyrate, and sulfalazine) exceeded solubility in the given volume of DMSO (5 µl). After confirming that these compounds still showed effect after a 10× dilution (as can be seen from hierarchical clustering in **Supplementary Figure S3**), the concentrations corresponding to the 1/10 highest oral dosages were used for these compounds. Detailed catalogue number and concentration of each compound is listed in **Supplementary Table S1**. After inoculation, the 96-well deep well plate was covered with a sterile silicone gel mat with a vent hole for each well made by a sterile syringe needle. Then, the plate was shaken at 500 rpm with a digital shaker (MS3, IKA, Germany) at 37°C for 24 hours in the anaerobic chamber.

### Metaproteomic sample processing and LC-MS/MS analysis

The sample processing was based on a previously reported metaproteomic sample processing workflow[25], we adapted it for microplates (**Supplementary Figure S1**). Briefly, after culturing for 24 hours, each 96-well plate was transferred out of the anaerobic station and was immediately centrifuged at 300 g for 5 min to remove debris. The supernatants were transferred into new 96-well deep well plates for another two rounds of debris removal at 300 g. The supernatants were then transferred to a new plate and centrifuged at 2,272 g for 1 hour to pellet the microbiome. The supernatant was removed and the pelleted bacterial cells were washed three times with cold PBS in the same 96-deep well plate, pelleting the cells after each wash by a 2,272 g centrifugation for 1 hour. The 96-well plate containing harvested bacterial cells was then stored overnight at −80°C before bacterial cell lysis and protein extraction. The lysis buffer was freshly prepared, containing 8 M urea in 100 mM Tris-HCl buffer (pH = 8.0), plus Roche PhosSTOP(™) and Roche cOmplete(™) Mini tablets. Microbial cell pellets were then re-suspended in 150 µl lysis buffer and lysed on ice using a sonicator (Q125 Qsonica, USA) with an 8-tip-horn probe. 100% amplitude was used (i.e. 15.6 Watts per sample), and four cycles of 30 s ultrasonication and 30 s cooling down were performed. Protein concentrations of the DMSO control samples were measured in triplicate using a detergent compatible (DC) assay (Bio-Rad, USA). Then, a volume equivalent to the average volume of 50 μg of protein in the DMSO control samples was acquired from each sample and placed into a new 96-deep well plate. The samples were reduced and alkylated with 10 mM dithiothreitol (DTT) and 20 mM iodoacetamide (IAA), followed by a 10× dilution using 100 mM Tris-HCl (pH = 8.0) and tryptic digestion under 37°C for 18 hours using 1 µg of trypsin per well (Worthington Biochemical Corp., Lakewood, NJ). Digested peptides were desalted using a panel of lab-made 96-channel filter tips generated by inserting 96 20 µl filter tips into a 96-well cover mat and stacking each filter tip with 5 mg of 10-μm C18 column beads. After being washed twice with 0.1% formic acid (v/v), tryptic peptides were then eluted with 80% acetonitrile (v/v)/0.1% formic acid (v/v).

After freeze-drying, each sample was re-dissolved in 100 μl 0.1% formic acid (v/v), and 2 μl of the solution (corresponding to 1 μg of proteins in the DMSO control) was loaded for LC-MS/MS analysis in a randomized order. An Agilent 1100 Capillary LC system (Agilent Technologies, San Jose, CA) and a Q Exactive mass spectrometer (ThermoFisher Scientific Inc.) were used. Peptides were separated with a tip column (75 μm inner diameter × 50 cm) packed with reverse phase beads (1.9 μm/120 Å ReproSil-Pur C18 resin, Dr. Maisch GmbH, Ammerbuch, Germany) following a 90 min gradient from 5 to 30% (v/v) acetonitrile at a 200 nL/min flow rate. 0.1% (v/v) formic acid in water was used as solvent A, and 0.1% FA in 80% acetonitrile was used as solvent B. The MS scan was performed from 300 to 1800 m/z, followed by data-dependent MS/MS scan of the 12 most intense ions, a dynamic exclusion repeat count of two, and repeat exclusion duration of 30 s.

### Assessment of the equal-volume strategy

Six dilutions of a single microbiome sample were prepared in triplicate wells and an equal volume was taken from each sample for tryptic digestion and LC-MS/MS analysis. Metaproteomic sample processing and analysis followed the same procedures stated above, and total peptide intensity was calculated. A DC protein concentration assay was also performed with each sample. Linearity between total protein concentration and total peptide intensity quantified by LC-MS/MS was then compared.

### Metaproteomics data analysis

#### 1) Metaproteomic database search

Protein/peptide identification and quantification, taxonomic assignment and functional annotations were done using the MetaLab software (version 1.1.0)[27]. MetaLab is a software that automates an iterative database search strategy, i.e. MetaPro-IQ [26]. The search was based on a human gut microbial gene catalog containing 9,878,647 sequences from http://meta.genomics.cn/. A spectral clustering strategy were used for database construction from all raw files, then the peptide and protein lists were generated by applying strict filtering based on a FDR of 0.01, and quantitative information of proteins were obtained with the maxLFQ algorithm on MaxQuant (version 1.5.3.30). Carbamidomethyl (C) was set as a fixed modification and oxidation (M) and N-terminal acetylation (Protein N-term) were set as variable modifications. Instrument resolution was set as “High-High”.

#### 2) Microbiome biomass analyses

Total microbiome biomass was estimated for each sample by summing peptide intensities. Taxonomic identification was achieved by assigning peptide sequences with lineage of lowest common ancestor (LCA). The “peptide to taxonomy” database (pep2tax database) was selected for mapping identified peptides to the taxonomic lineages [27]. Bacteria, eukaryota, viruses, and archaea were included in the LCA calculation. Taxonomic biomass was quantified by summing the intensities of the peptides corresponding to each taxon. A Bray-Curtis dissimilarity-based approach [28] was applied for evaluating the variation of genus-level biomass contributions between drug-treated and DMSO control groups. Calculation of the Bray-Curtis distance was performed using the R package “vegan”[29].

#### 3) Functional analysis

The quantified protein groups were first filtered according to the criteria that the protein appears in >80% of the microbiomes with at least one drug treatment. Then LFQ protein group intensities of the filtered file was log_2_-transformed and normalized through quotient transformation (x/mean) using the R package clusterSim. Then, LFQ protein group intensities were processed by a ComBat process [54, 55] using iMetalab.ca [56] to remove possible batch effects between individual microbiomes. Using the ComBat-corrected data, an unsupervised non-linear dimensionality reduction algorithm, t-distributed stochastic neighbor embedding (t-SNE)[30] was then applied to visualize similarities between samples using the R package Rtsne. Parameter for the function Rtsne() were, perplexity=10, max_iter = 1200 (number of iterations), other parameters were set as default. The R function geom_polygon implemented in ggplot2 was used to visualize the t-SNE results.

Functional annotations of protein groups, including COG and KEGG information, were obtained in the MetaLab software. In addition, KEGG ortholog (KO) annotation of protein FASTA sequences was conducted using GhostKOALA (https://www.kegg.jp/ghostkoala/)[57]. Log_2_ fold-change of each drug-treated sample relative to the corresponding DMSO control was calculated using the abundances of proteins annotated to COG categories and COGs. Functional enrichment analysis was performed using the enrichment module on iMetalab.ca through inputting the list of COG functional proteins. Adjusted *p*-value cutoff was set at 0.05 for the enrichment analysis.

### Statistical analysis

We examined data distribution on all levels of data, and results indicated non-normal distributions of the dataset (examples shown in **Supplementary Figures S10 and 11**). Hence, a non-parametric statistical hypothesis test, the Wilcoxon rank sum test, was applied in statistical analyses. For multiple comparisons, *p*-values were adjusted using the Benjamini-Hochberg false discovery rate (FDR) procedure[58]. For multivariate analysis, partial least-squares discriminant analyses (PLS-DA) based on ComBat-corrected protein group intensities were performed using MetaboAnalyst (http://www.metaboanalyst.ca/)[59]. PLS-DA model were evaluated by cross-validation of R^2^ and Q^2^.

### Data visualizations

Box plots, violin plots, hierarchical clustering, 3D scatter plots, heatmaps, PCA, and t-SNE were visualized using R packages ggplot2, gridExtra, scatterplot3d, and pheatmap. Pathway maps were visualized using iPATH 3 (https://pathways.embl.de/)[60] and Pathview Web (https://pathview.uncc.edu/)[61]. Stacked column bars and functional enrichments were visualized on iMetaLab.ca.

## List of abbreviations

COG: Clusters of orthologous groups
DMSO: Dimethyl sulfoxide
FDR: False-discovery rate
FOS: Fructooligosaccharide
GABA: Gamma-Aminobutyric Acid
KEGG: Kyoto Encyclopedia of Genes and Genomes
LC-MS/MS: Liquid chromatography–tandem mass spectrometry
LFQ: Label-free quantification
NSAIDs: Nonsteroidal anti-inflammatory drugs
PCA: Principle component analysis
PLS-DA: Partial least squares discriminant analysis
POC: Proof-of-concept
VIP: Variable importance in projection

## Declarations

### Ethics approval and consent to participate

The Research Ethics Board protocol (# 20160585-01H) for stool sample collection was approved by the Ottawa Health Science Network Research Ethics Board at the Ottawa Hospital. All volunteers signed consent forms.

### Consent for publication

Not applicable.

### Competing interests

D.F. and A.S. have co-founded Biotagenics and MedBiome, clinical microbiomics companies. All other authors declare no competing interests.

### Funding

This work was supported by the Government of Canada through Genome Canada and the Ontario Genomics Institute (OGI-114 and OGI-149), CIHR grant (ECD-144627), the Natural Sciences and Engineering Research Council of Canada (NSERC, grant no. 210034), and the Ontario Ministry of Economic Development and Innovation (REG1-4450).

### Author contributions

D.F., A.S. and L.L. designed the study. J.M. coordinated the volunteers and collected the samples. L.L., X.Z. and Z.N. developed the workflow and performed the experiments. L.L., Z.N., X.Z. and K.C. analyzed and visualized the data. L.L, D.F. and A.S. wrote the manuscript. All authors read and approved the final manuscript.

## Acknowledgments

The authors wish to thank Dr. Kendra Hodgkinson for editing the manuscript.

## Supplementary figures

**Figure S1.**
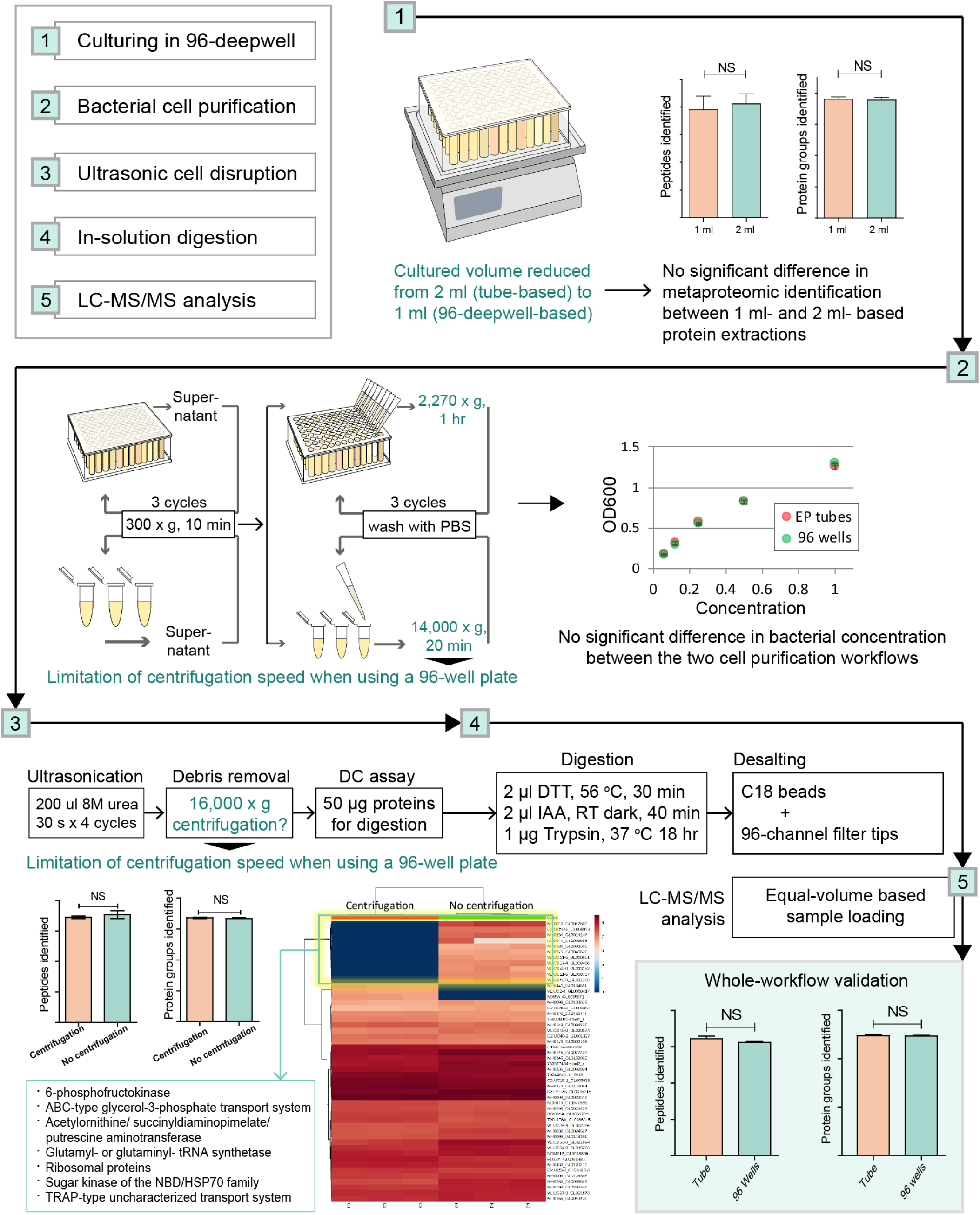
Establishment and step-by-step validation of the microplate-based metaproteomic sample preparation workflow of the RapidAIM assay. After culturing in a 96-well deepwell plate [1], the bacterial cells were washed with PBS [2], then disrupted with four cycles of 30 s ultrasonication [3]. Protein concentration was measured in the DMSO control sample using a DC protein assay. This concentration was used to calculate a volume equivalent to 50 µg proteins in the control sample. This volume was taken from every sample and digested with trypsin [4], then desalted with a panel of 96 filter tips packed with C18 beads and equal volumes of each sample were analyzed by LC-MS/MS [5]. Due to a few differences in the metaproteomic procedure compared with tube-based protocols, we examined the effect of protocol differences on sample quality step-by-step. In previously published protocols, samples were cultured in 2 ml culture tubes^1,2^. In step [1], culturing samples in a 96-well plate reduced the sample size to 1 ml. No significant difference was shown by *t*-test in peptide and protein group identification numbers between protein extractions from 1 ml and 2 ml samples. For step [2], the 96-deepwell plates limited centrifugation speed to 2,270 g (versus 14,000 g in the original protocol), so we extended the centrifugation time from 20 min to 1 hour and tested whether bacterial concentration was affected by the altered centrifugation process^1,3^. Concentration of purified bacterial cells were compared by OD_600_ after being re-suspended in 1 ml, 2 ml, 4 ml, 8 ml, and 16 ml PBS. No significant difference in OD_600_ reads was observed between the two cell purification protocols. In step [3], the original protocol used a high-speed centrifugation (16,000 g) to remove cell debris after the ultrasonication. Due to the limitation of centrifugation speed when using a microplate, we compared metaproteomic profiles of the sample when digested with or without cell debris removal. No significant differences were found in the number of protein identifications. In terms of differentially identified proteins, we found that the samples without cell debris removal resulted in more identifications of cell-membrane proteins such as the ABC-type transport systems, as well as cytoskeleton-related proteins, such as the translation elongation factor EF-Tu^4,5^, 6-phosphofructokinase^6^, and ribosomes^7^. Therefore, we infer that eliminating the centrifugation process could reduce the removal of cytoskeleton and cell membrane proteins. For step [4], no validation was necessary as the same type of filter tips were used in both protocols. Finally, we performed a whole-workflow comparison of the metaproteomic outcomes between traditional tube-based and 96-well-based processes. The microplate-based metaproteomic workflow showed no statistically significant difference in peptide and protein identification compared with the tube-based workflow.

**Figure S2.**
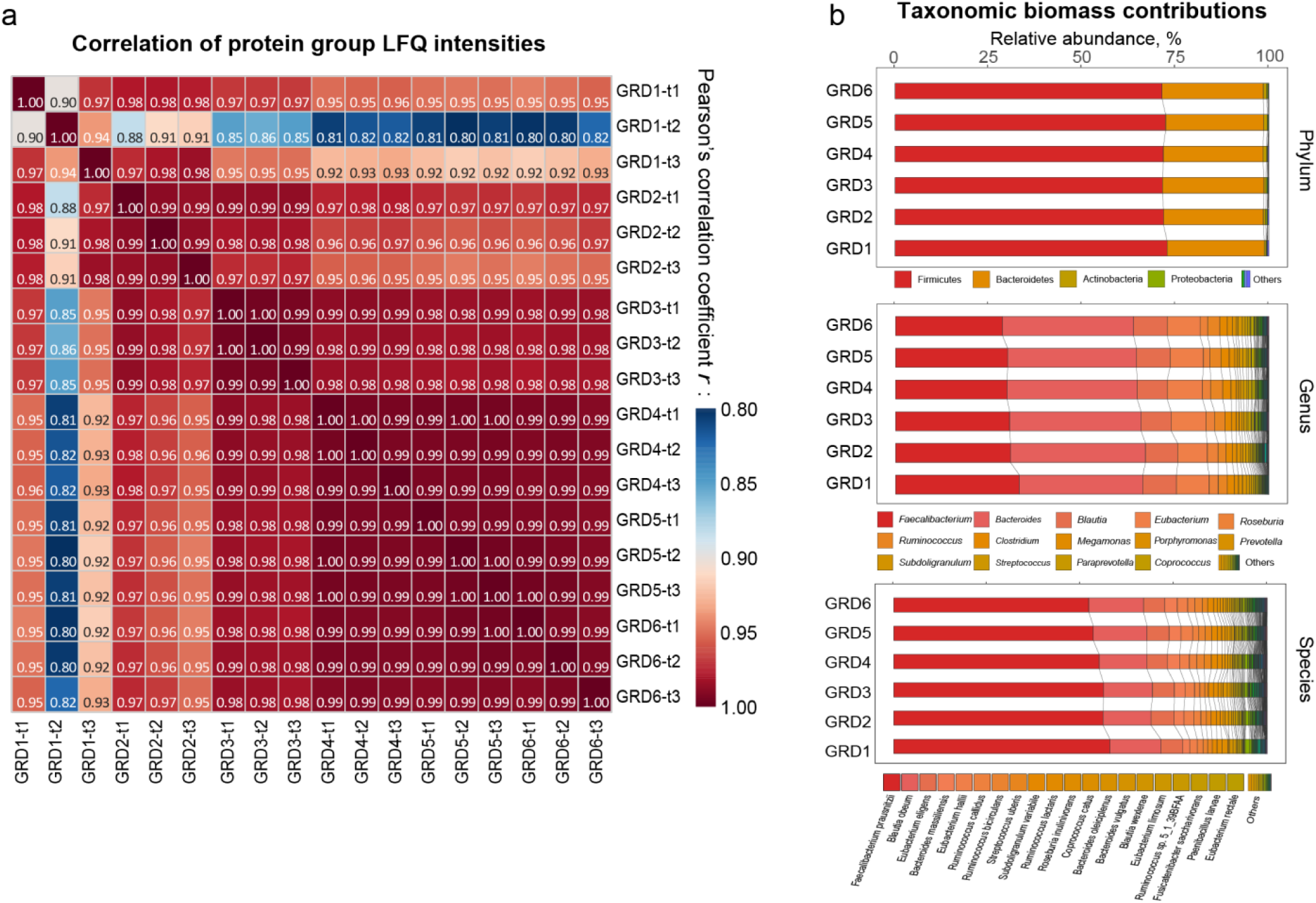
Assessment of the equal-volume digestion and LC-MS/MS analysis strategy. (**a**) In triplicates, six dilutions of a single microbiome sample were subjected to this equal-volume based analysis. The LFQ intensities of protein groups showed Pearson’s correlation coefficient *r* > 0.95 between most dilutions, but a lower *r* was seen in the samples with the lowest concentration. (**b**) Comparison of taxonomic biomass contributions on different levels (summed peptide intensity assigned to different taxa) among diluted groups suggested very low level of bias. GRD1-6 are six different dilution gradients (protein concentration shown in Figure 1b), and t1-t3 are technical replicates of the same conditions.

**Figure S3.**
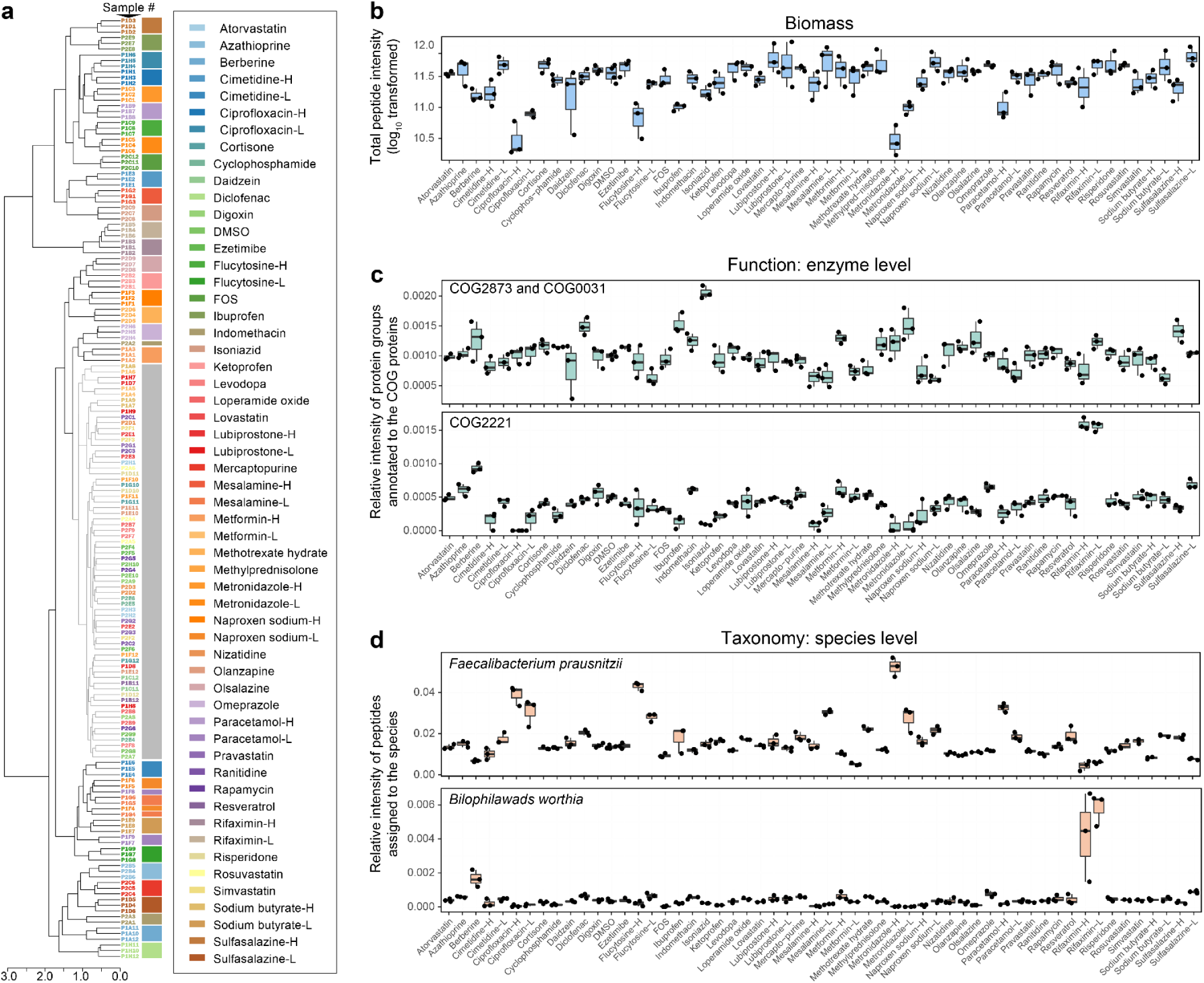
Reproducibility of RapidAIM assay on different levels. (**a**) Clustering using protein group composition information showed that triplicates of drug treatments were closely clustered. A hierarchical tree was generated based on Pearson’s correlation coefficient. The cluster corresponding to the gray box contained DMSO control samples; samples in this cluster indicated drugs that had very weak effects on the microbiome. (**b**) Box chart suggesting that biomass effects by drugs are highly reproducible. (**c**) Examples showing reproducible functional responses to drugs at the enzyme level. COC0031 and COG2873 are cysteine synthase and O-acetylhomoserine/O-acetylserine sulfhydrylase (pyridoxal phosphate-dependent), respectively; both enzymes are involved in the conversion of sulfide to L-cysteine^8^. COG2221 is dissimilatory sulfite reductase (desulfoviridin), alpha and beta subunits; this enzyme reduces sulfite to sulfide^9^. (**d**) Examples showing reproducible taxonomic responses to drugs at the species level. *F. prausnitzii* is a ubiquitous bacterium of the intestinal microbiota^10^; *B. worthia* is a taurine-degrading bacterium which can reduce sulfite to sulfide by dissimilatory sulfite reductase^9^. COG2221 and *B. worthia* show a correlation in their response to different drugs. Box spans interquartile range (25th to 75th percentile), and line within box denotes median.

**Figure S4.**
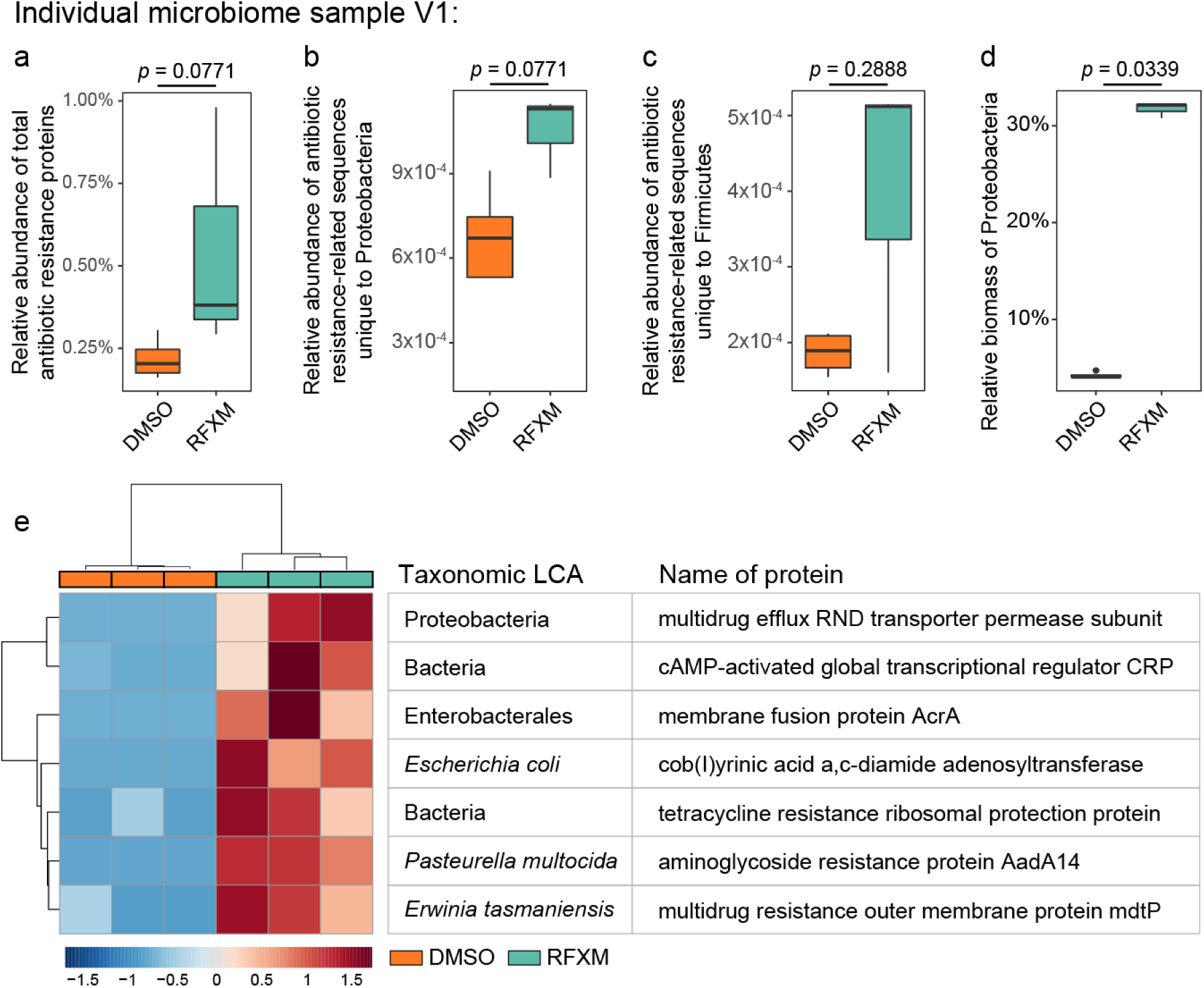
Case study on microbiome V1’s response to rifaximin (RFXM). We have shown that the antibiotic rifaximin did not show overall biomass inhibition on microbiome sample V1 (Figure 2a and Supplementary Figure S3b). We examined whether the expressions of antibiotic resistance proteins were affected. Briefly, the LC-MS/MS raw files were specifically searched against the Structured Antibiotic Resistance Genes (SARG) database 48 which highlighted 118 antibiotic resistance protein groups across the dataset. (**a**) We found an increase in relative abundance of total antibiotic resistance proteins in microbiome V1 in response to rifaximin. (**b and c**) Particularly, antibiotic resistance peptide sequences unique to Proteobacteria and Firmicutes were increased. (**d**) Moreover, despite no significant change in total microbiome biomass in V1, a significant 6.5-fold increase in the relative biomass of Proteobacteria in the whole microbial community was observed. (**e**) Non-parametric test resulted in seven significantly increased antibiotic resistance protein groups (FDR-adjusted p value<0.05). These protein groups belonged predominantly to Proteobacteria (5 out of 7). Increase of Proteobacteria is associated with dysbiosis in gut microbiota 49. These together suggested a potential risk of rifaximin administration in individual V1. (*p* values were based on two-sided Wilcoxon test; box spans interquartile range (25th to 75th percentile), and line within box denotes median.

**Figure S5.**
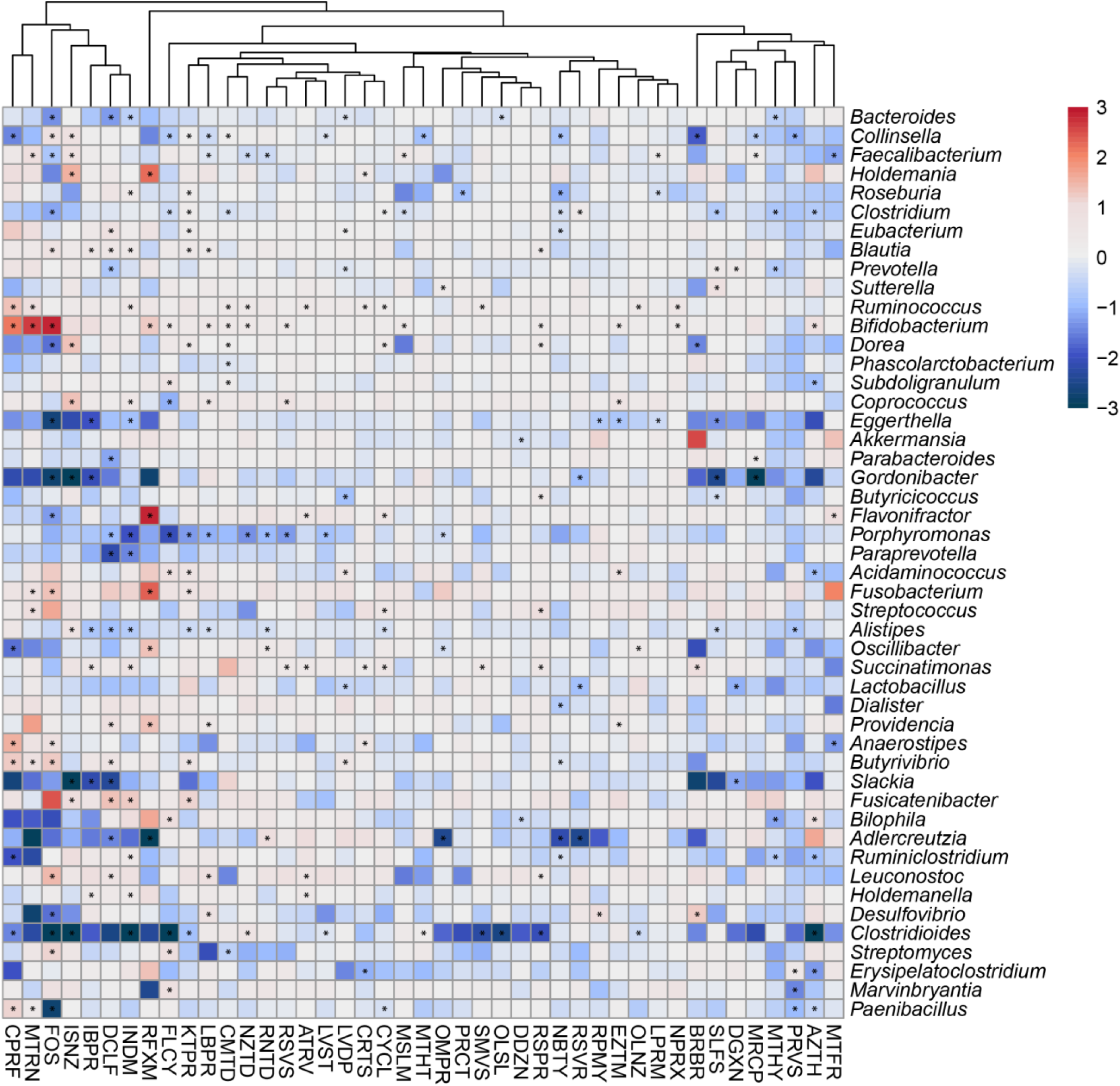
Log2 fold-change of relative abundance at the genus level in response to each drug compared with the DMSO control. Genera that existed in ≥80% of the volunteers are shown. Star (*) indicate significantly changed bacterial abundance by Wilcoxon test, *p* <0.05..

**Figure S6.**
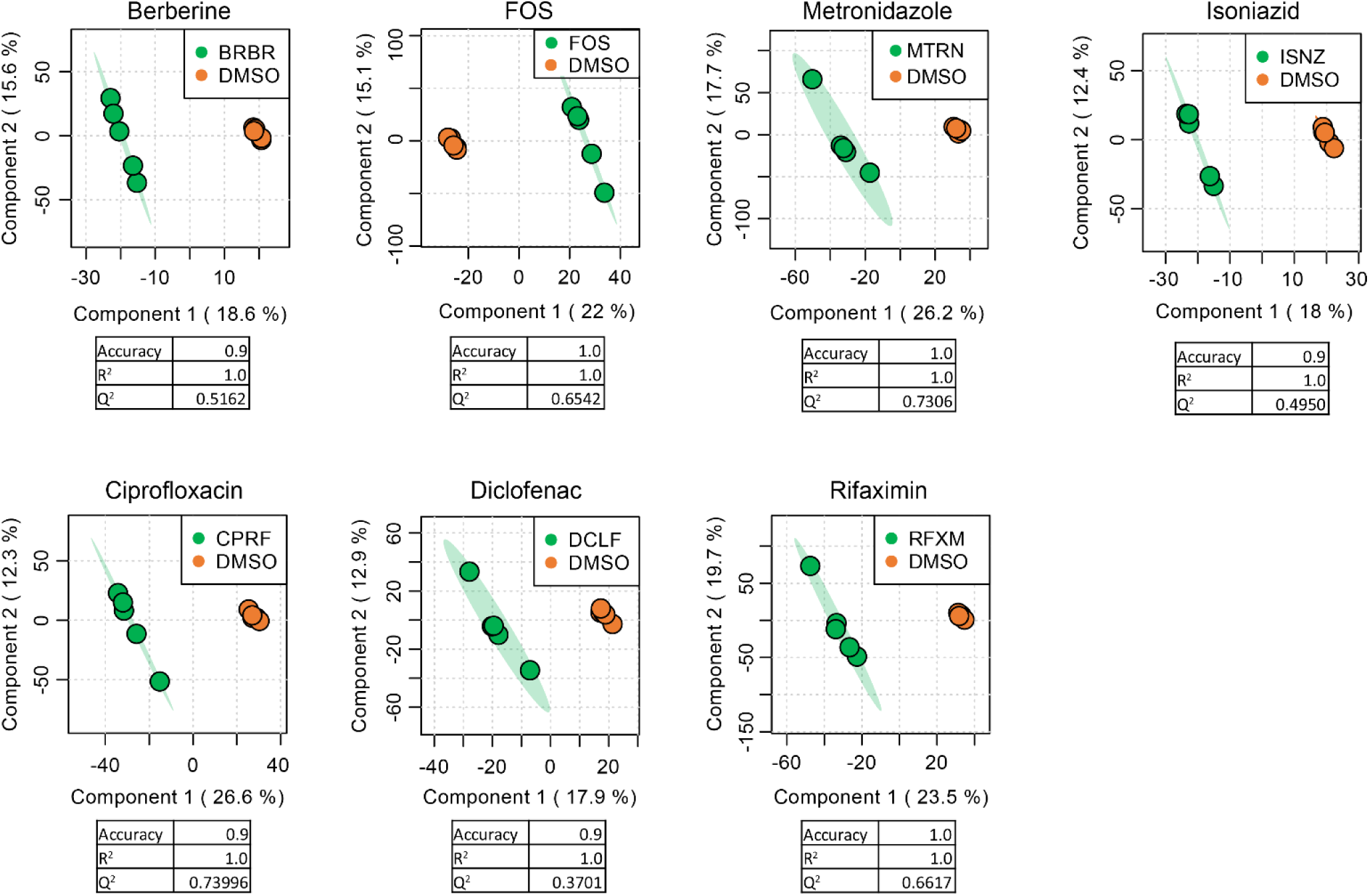
Score plots and cross-validations of seven PLS-DA models. PLS-DA models of microbiomes responses to each compound were established using MetaboAnalyst 4.0. PLS-DA model qualities were assessed through cross-validation, and accuracy, R^2^ and Q^2^ were given for each model. Seven compounds were found with valid PLS-DA models distinguishing the effect of the compound from the DMSO control.

**Figure S7.**
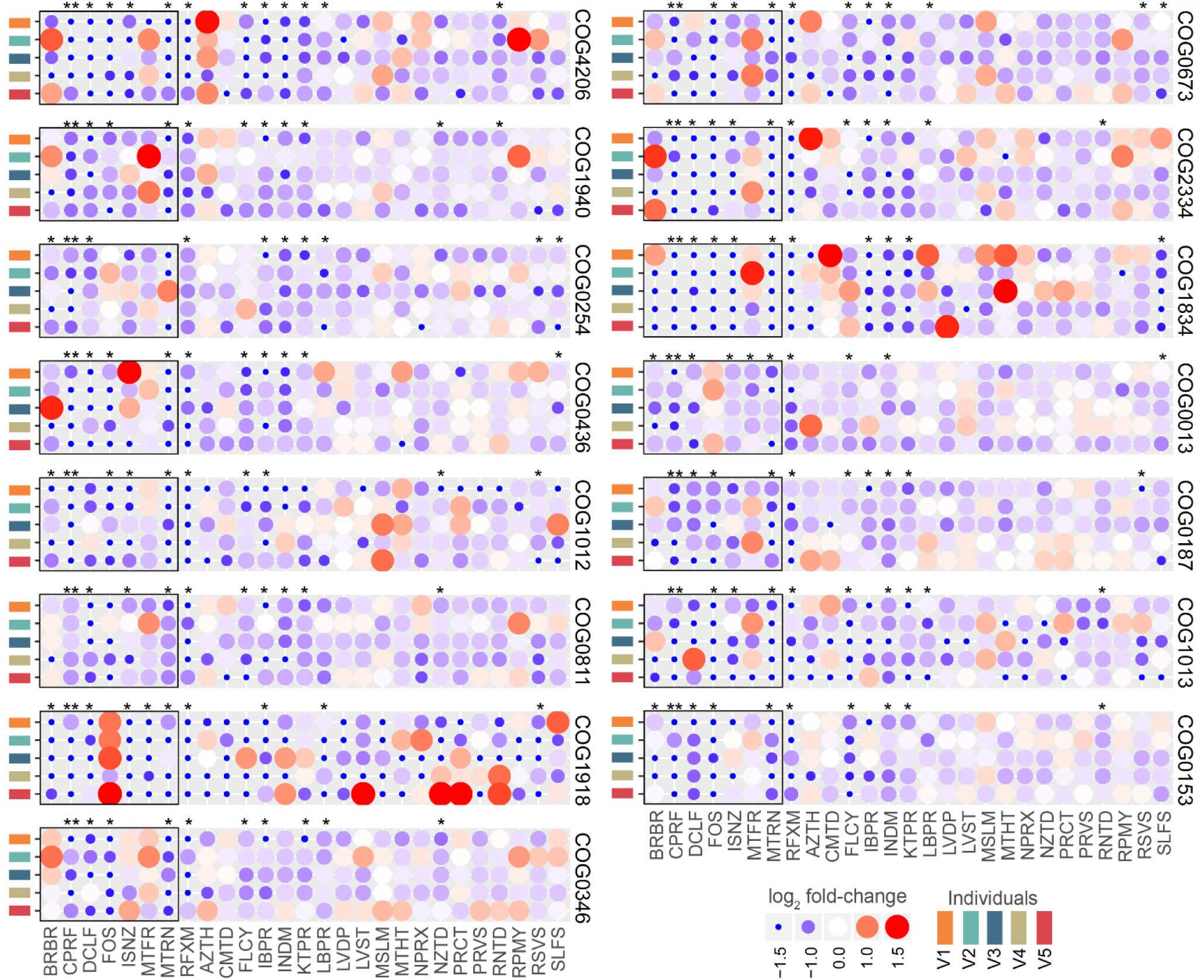
Log_2_ fold-change of functions at the COG protein level. On the COG functional protein level, 535 COGs were significantly decreased by at least one drug treatment. The 15 COGs that were affected by ⩾ 10 compounds are shown in this figure. Statistical significance was evaluated by one-sided Wilcoxon rank sum test, FDR-adjusted *p*-values: *, *p* < 0.05; **, *p* < 0.01.

**Figure S8.**
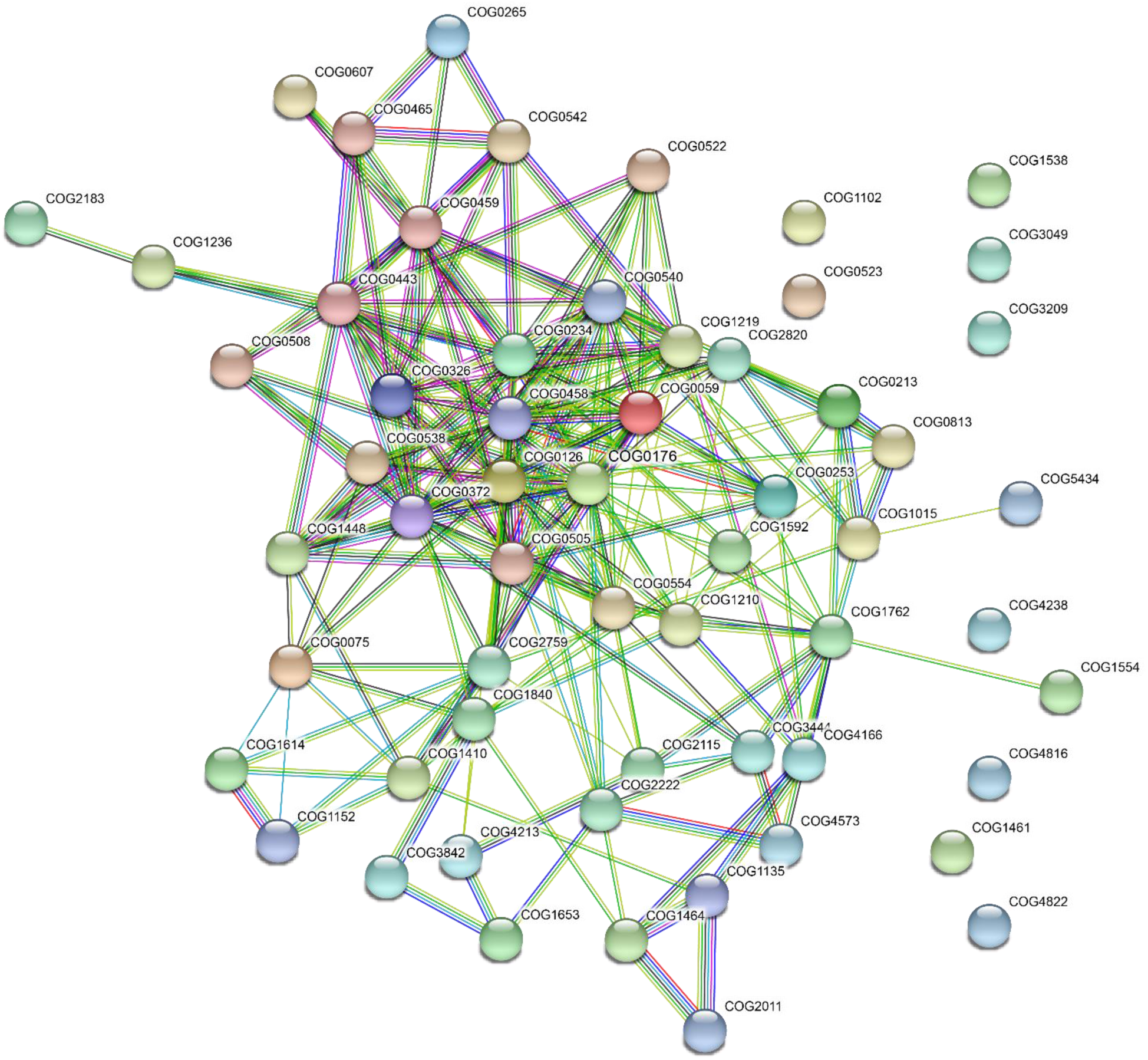
String interaction of COG functional proteins significantly stimulated by diclofenac.

**Figure S9.**
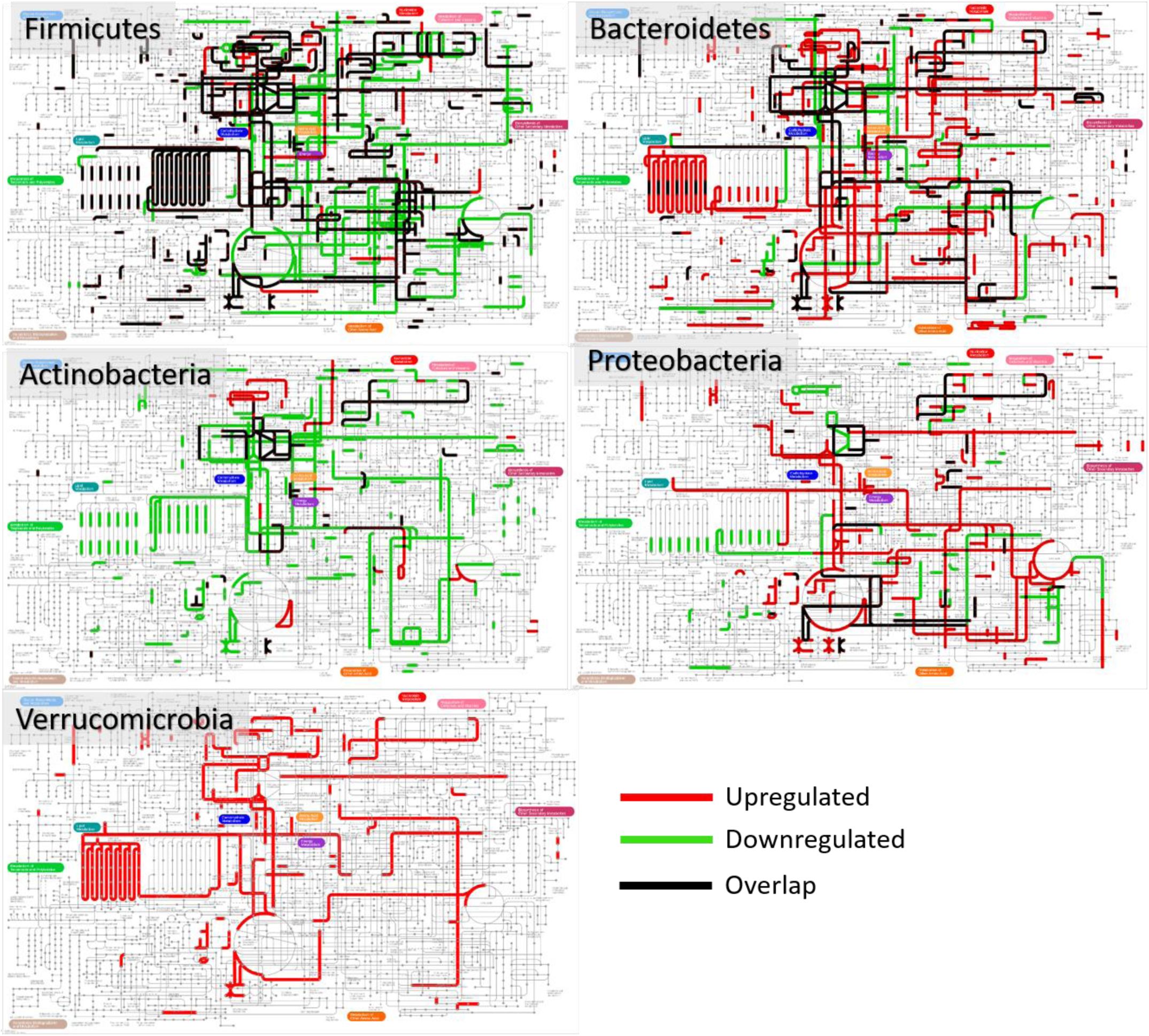
Phylum-specific functional responses to Berberine. Protein groups with VIP scores of >1 were annotated to phyla and COGs. Up- and down-regulated (red and green lines) COGs corresponding to different phyla were illustrated on a metabolic pathway map using iPath 3. Pathway maps for each phylum were combined, and overlapped pathways were shown in black lines. Our data suggest that different phyla responded differently at a functional pathway level.

**Figure S10.**
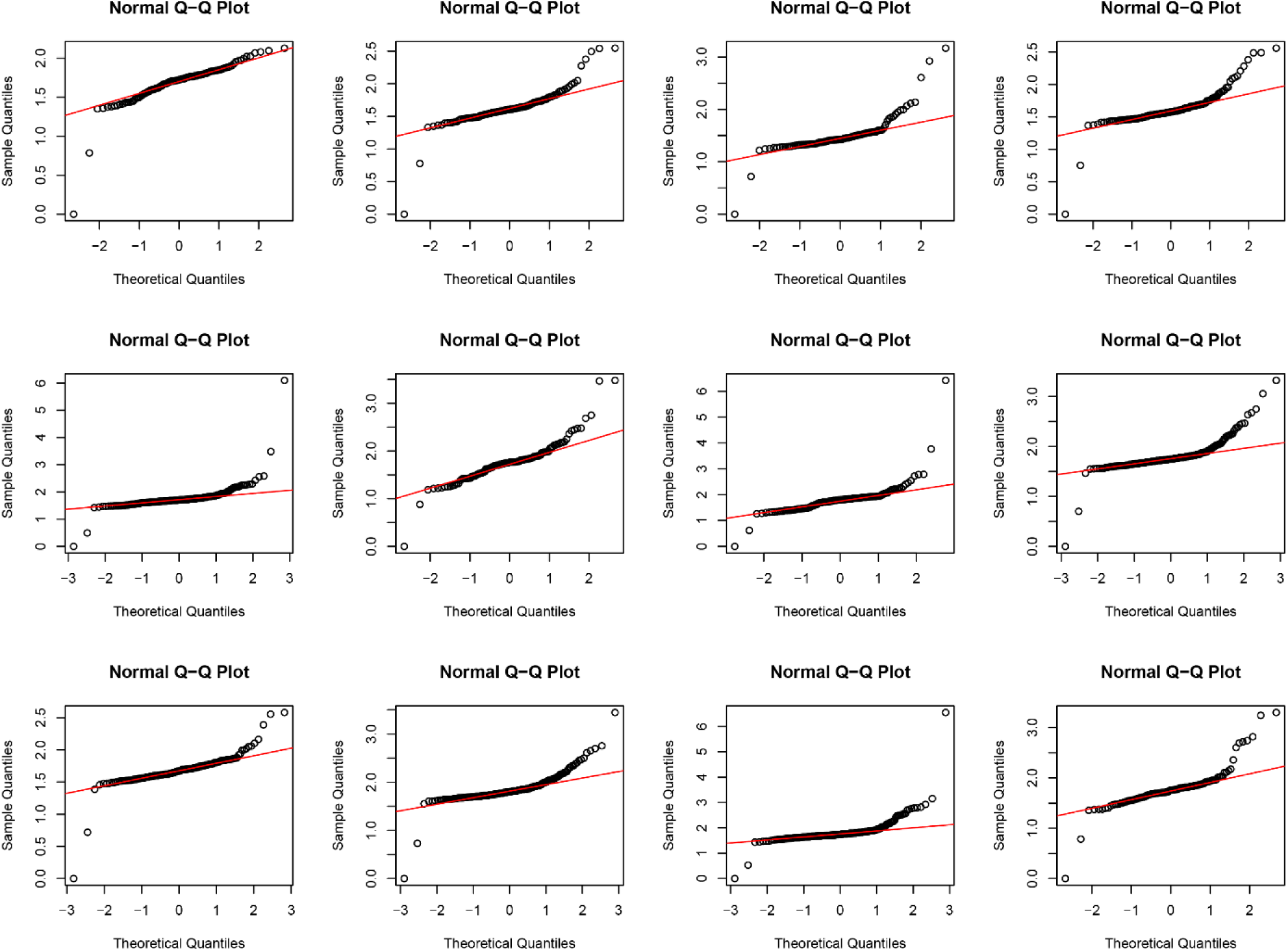
Randomly selected LFQ intensities of protein groups showing heavy tailed distribution on the Q-Q plots.

**Figure S11.**
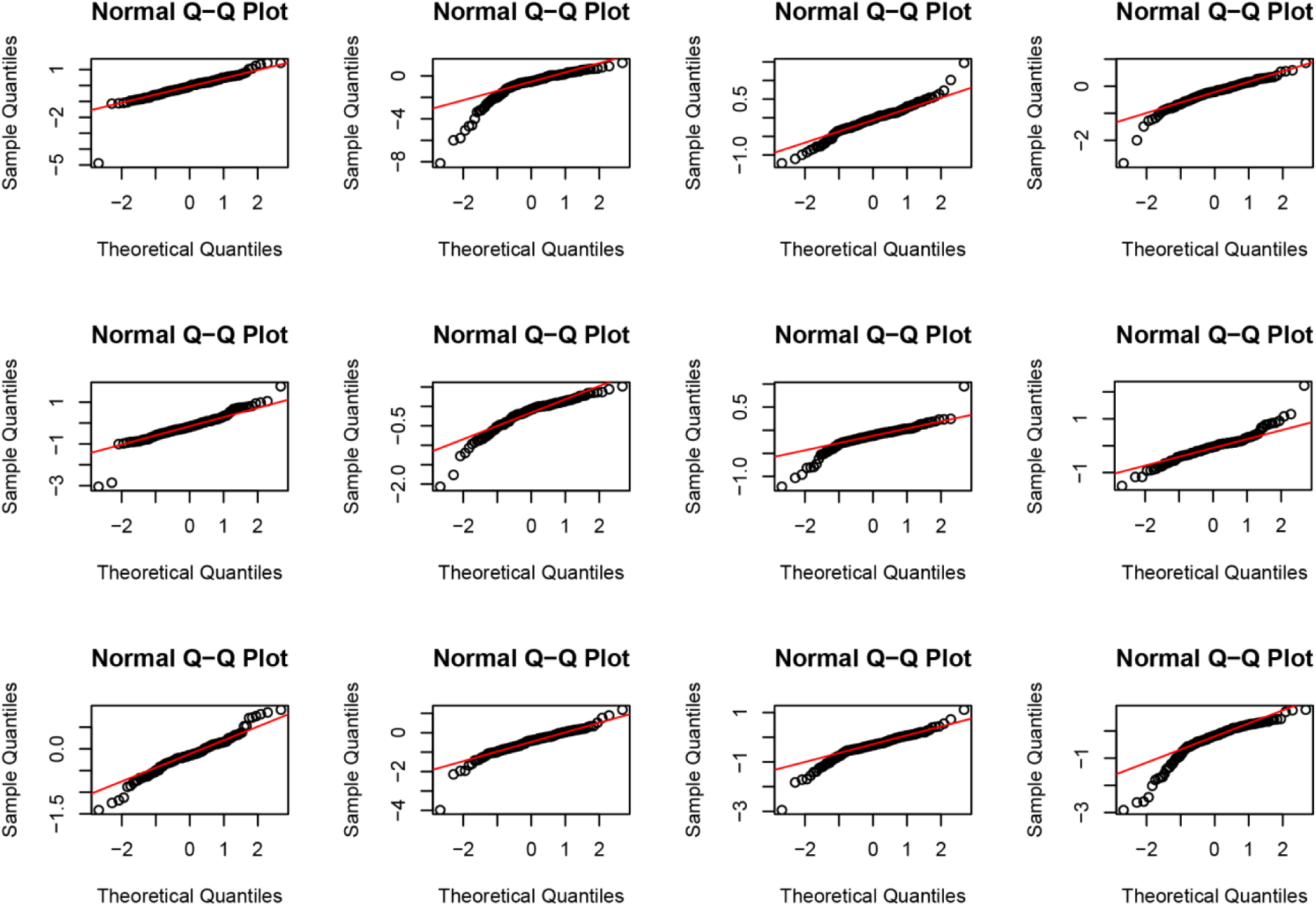
Randomly selected log_2_-fold changes of COGs showing heavy tailed distribution on the Q-Q plots.

